# The Temporal Dynamics of Metacognitive Experiences Track Rational Adaptations in Task Performance

**DOI:** 10.1101/2023.09.26.559523

**Authors:** Luc Vermeylen, Senne Braem, Ivan I. Ivanchei, Kobe Desender, J.M. García-Román, Carlos González-García, María Ruz, Wim Notebaert

**Author notes:** **Corresponding Author:** Luc Vermeylen Department of Experimental Psychology Ghent University Henri Dunantlaan 2, 9000 Ghent Belgium.

## Abstract

Human task performance elicits diverse subjective metacognitive experiences, such as boredom, effort, fatigue and frustration, which are thought to play important roles in the monitoring and regulation of cognitive processes. Yet, their specific contributions to task performance remain poorly understood. Therefore, we investigated the temporal dynamics underlying these metacognitive experiences and the latent cognitive processes supporting task performance. We used a time-on-task design using a conflict Flanker task, and analyzed the data using a comprehensive approach encompassing behavioral, model-based, subjective, and neural measures. Our results show that the temporal dynamics in cognitive processes can be understood as a rational attempt to optimize task performance and that distinct metacognitive experiences track different aspects of this rational endeavor. These findings suggest that metacognitive experiences act as tools for humans to gain insights into the optimality of their cognitive performance.

## Introduction

Humans tend to have several subjective metacognitive experiences when performing cognitive tasks^1,2^. For example, we often have an internal sense of certainty or confidence with respect to our performance – a well-studied feature of decision-making that facilitates performance in the absence of feedback^3–6^. However, task performance tends to be accompanied by a much broader spectrum of metacognitive experiences, including feelings of boredom, effort, fatigue and frustration. Consider, for example, a young student immersed in an examination setting: the challenging nature of the exam may elicit a sense of frustration, prompting the student to pause, reevaluate, and adopt a more cautious strategy. Hence, these experiences are thought to provide valuable insights in task performance and serve as a guide for strategic adjustments^3–7^. However, the precise relationship between such experiences and cognitive task mechanisms remains unclear. Nevertheless, a deeper understanding of how (and which) metacognitive experiences influence task performance could pave the way for targeted training programs aimed at enhancing awareness of the experiences that exert the most substantial impact on performance^8,9^.

One of the common challenges in maintaining optimal levels of task performance is the presence of irrelevant stimuli in the environment that capture our attention away from the primary task. This phenomenon, which we will refer to as irrelevant capture, emphasizes the importance of monitoring our environment and the need for behavioral regulation. For instance, imagine the young student distracted by subtle hallway noises, a flickering light, or a ticking clock. To regain his focus and align his behavior with his academic goal, he first needs to register the distraction and then take steps to restore his concentration. This ability to adapt our information processing system in response to environmental changes falls under the domain of cognitive control and encompasses the mental processes essential for managing goal-directed behavior in the presence of irrelevant stimuli or conflict, where mutually incompatible representations compete for control over action. While traditional accounts described conflict as the trigger for behavioral adaptations^8^, more recent accounts have argued that the negative metacognitive experiences associated with cognitive control and conflict might be the trigger instead^9–11^. These experiences include aversive reactions towards the conflicting stimuli themselves (what we will refer to as “conflict aversiveness”^12^), but also other feelings of effort, boredom, frustration and fatigue^13,14^. Therefore, contemporary accounts of cognitive control take into account that there are experiential costs (and benefits) associated with investing cognitive control, and propose that these subjective experiences may serve as influential forces driving behavioral regulation^9,10,15–17^.

While it has become evident that cognitive control and conflict elicit a broad palette of negative metacognitive experiences, the precise experiences that offer metacognitive insights into the underlying cognitive processes of task performance remain unclear. Equally uncertain is the manner in which these experiences act as sources for making adaptive changes to guide and optimize performance. While attempts are being made to construct normative computational accounts of how such subjective experiences could arise and what their exact signaling and regulatory function could be^18^, there is a scarcity of empirical data that simultaneously tracks the evolution of task performance and the evolution of multiple subjective experiences over time, which is crucial for advancing theoretical understanding. The dimension of time is important here, as these metacognitive experiences and task performance are not static but constantly evolving. Therefore, in the present study we aim to address this gap by investigating the temporal dynamics between multiple metacognitive experiences and the cognitive processes underpinning task performance such as the setting of one’s decision boundary or susceptibility to irrelevant capture. Specifically, we investigated which type of metacognitive experiences were related to changes in well-studied parameters of cognitive task performance. Across two studies (behavioral study, N = 66; EEG study, N = 45), human participants engaged in a standard conflict Flanker task across 18 blocks of time-on-task, spanning nearly two hours. After each block, participants reported their metacognitive experiences of conflict aversiveness, boredom, effort, fatigue and frustration (Fig. 1).

**Fig 1.**
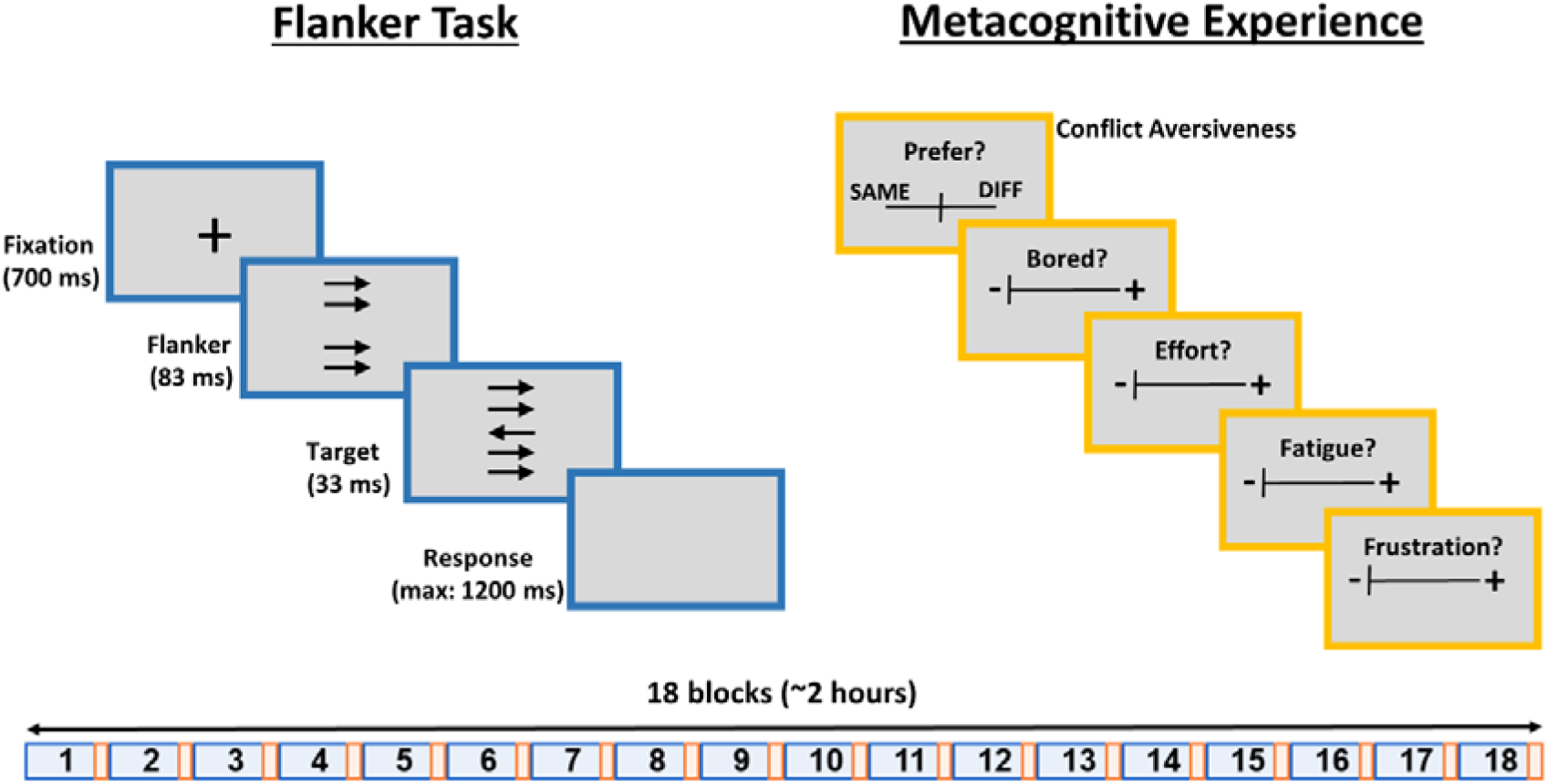
Task design. Participants performed a Flanker task (adapted from Fischer et al., 2018) for 18 blocks, which lasted roughly two hours. After each block, participants reported on their metacognitive experiences (conflict aversiveness, boredom, effort, fatigue, frustration) using visual analogue scales. To measure conflict aversiveness, participants reported whether they preferred being presented congruent (“SAME”) or incongruent (“DIFF”) trials by moving the marker away from the middle (which is the indifference point, i.e., no preference). For the other measures of metacognitive experience, the scale always started at the left-most position, denoting that the experience was not present at all (whereas the right-most position denoted that the experience was highly present).

Given the intricate challenge of quantifying the dynamic temporal interplay between these two domains, we relied on a time-on-task design which we analyzed using a comprehensive approach encompassing behavioral, subjective, and neural measures, as well as several theory-driven computational and analytic approaches (i.e., sequential sampling modeling, Bayesian multivariate modeling, representational similarity analysis, normative modeling). As a result, our approach allowed us to explore the dynamic interplay between metacognitive experiences and task performance with a high level of detail.

To set the stage for the results to follow, we started by dissecting the temporal dynamics of behavioral task performance in model-based cognitive mechanisms using a sequential sampling model, which we subsequently verified as a biologically plausible account of decision formation. Next, we assessed the temporal dynamics of metacognitive experiences and quantified how they were related to the temporal dynamics of the task performance parameters. Finally, we show that the main task performance dynamics can be understood as rational attempts to optimize an objective criterion (i.e., reward rate) and that distinct metacognitive experiences track distinct cognitive operations to achieve that goal. Together, these findings suggest an important role for metacognitive experiences in relation to strategic adaptations during task performance and offer specific pathways through which they may influence these processes.

## Results

### Diffusion modeling of task performance

To increase power, we collapsed the sample across the behavioral and EEG experiment where possible (N = 111). We observed clear behavioral changes with time-on-task (Fig. 2A). Participants became faster while maintaining a stable error rate and suffered less irrelevant capture (i.e., reductions in both reaction time [RT] and error rate congruency effects, see Supplementary Appendix S1 for details). While previous time-on-task studies typically relied on analyses of average RTs and error rates (e.g.^19–21^), sequential sampling models of decision formation, such as the drift diffusion model (DDM), allow us to model task performance in terms of theoretically grounded, biologically plausible, cognitive mechanisms^22^. In the DDM, decisions are formed through the accumulation of evidence over time which is governed by three parameters: First, the rate or efficiency of the accumulation is determined by the drift rate. Second, accumulation stops at a certain decision boundary, which quantifies one’s level of cautiousness and accounts for the speed-accuracy tradeoff. Third, additional processes that contribute to the RT outside the decision itself are captured by the non-decision time.

**Fig 2.**
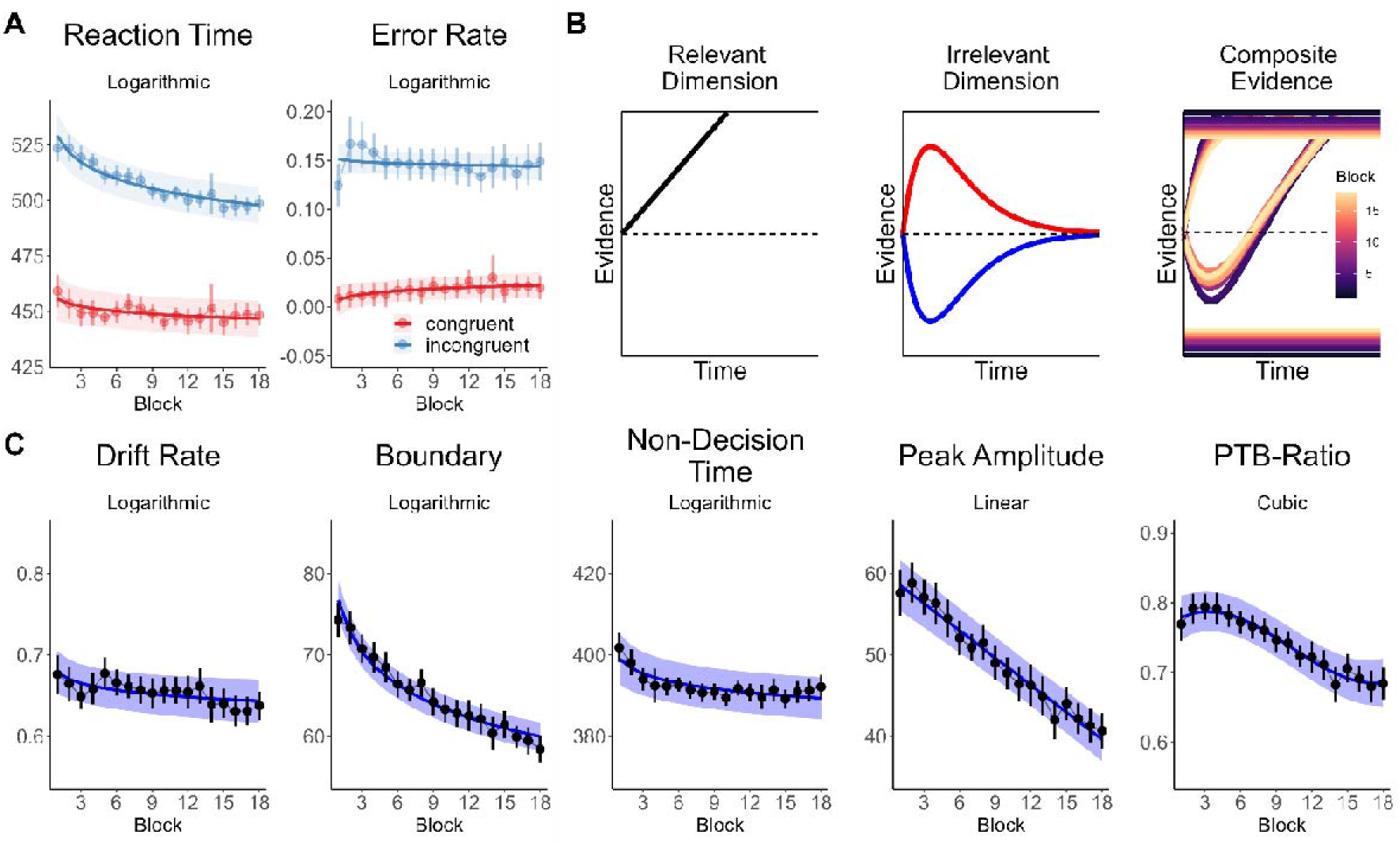
Task performance. **(A)** Time-on-task effects in behavior (RT and accuracy). **(B)** Functional form of the latent decision variable for the relevant (constant drift rate) and irrelevant dimension (time-varying drift rate following a gamma impulse function with its main parameter peak amplitude) and evolution of the composite latent decision variable and decision boundary over time-on-task. The upper boundary represents the correct response while the lower boundary represents the error response. Red denotes activation of the irrelevant dimension on congruent trials while blue denotes activation of incongruent trials. **(C)** Time-on-task effects on the DDM parameters. The subtitle denotes the best model for time-on-task (linear, logarithmic, quadratic or cubic) based on BIC. PTB-Ratio = peak-to-bound ratio. Error bars represent the 95% within-subject CI.

However, the standard DDM fails to fully account for the patterns observed in RT distributions of conflict tasks^23^. To account for such patterns, instead of capturing the process of evidence accumulation with a single parameter, it can be further decomposed into contributions from the relevant and irrelevant stimulus dimensions^24^. The relevant dimension, such as the target in a Flanker task, is assumed to have a constant drift rate. On the other hand, the irrelevant dimension, such as the distractors in a Flanker task, has a time-varying drift rate modeled using a gamma impulse response which varies in terms of its peak amplitude (Fig. 2B). The latter parameter represents how much the irrelevant dimension influences decision formation in a short-lived manner, i.e., the degree of “irrelevant capture” or the performance costs due to conflict (often termed “interference effects” in the context of RT or accuracy measures).

To investigate the effects of time-on-task, we fitted this extended DDM to the behavioral data of each block and obtained four parameter estimates per block. Subsequently, we examined the impact of time-on-task on these parameters by fitting separate Linear Mixed Effect Models for linear, logarithmic, quadratic, or cubic effects of time-on-task. We chose the model with the lowest Bayesian Information Criterion (BIC, see Methods).

The results revealed slight logarithmic decreases in drift rate, β (standardized Beta) = - 0.058, *SE* (standard error) = 0.023, *t*(110) = -2.57, *P* = .012, and non-decision time, β = - 0.084, *SE* = 0.022, *t*(110) = -3.80, *P* < .001, showing that the rate of evidence accumulation decreased, and less time was spent on processes outside decision formation over time (Fig. 2C). Decision boundary showed a strong logarithmic decrease, β = -0.325, *SE* = 0.023, *t*(110) = -13.88, *P* < .001, suggesting participants became less cautious. Furthermore, peak amplitude exhibited a strong linear decrease, β = -0.300, *SE* = 0.023, *t*(110) = -13.09, *P* < .001, implying that evidence from the irrelevant dimension had a reduced impact over time.

However, given that the decision boundary changes with time-on-task, peak amplitude is no longer a pure measure of irrelevant capture, as its influence on decision formation depends on the height of the decision boundary. For instance, a peak amplitude value of 20 evidence units indicates more interference when the boundary is set at 30 compared to when it is set at 60. Therefore, to provide a more reliable assessment of irrelevant capture, it is crucial to consider changes in the decision boundary alongside peak amplitude. Thus, we introduced a novel metric, the peak-to-bound ratio (PTB-ratio). By normalizing peak amplitude by the decision boundary (dividing peak amplitude by the boundary), we obtain a measure that reflects the impact of the irrelevant dimension independently of the decision boundary setting. Consequently, this approach ensures that the parameters uniquely align with their proposed underlying mechanisms. The peak-to-bound ratio showed a cubic pattern: an early increase followed by a decrease, plateauing towards the end (β*_linear_* = -0.305, *SE* = 0.037, *t*(427.33) = - 8.20, *P* < .001; β*_quadratic_* = -0.022, *SE* = 0.015, *t*(1774) = -1.90, *P* = .058; β*_cubic_* = -0.111, *SE* = 0.029, *t*(1774) = 3.85, *P* < .001).

Together, these diffusion modeling results show that participants mainly became less cautious while the degree of irrelevant capture decreased. For more details on the diffusion model, fit assessment and parameter recovery, see Methods and Supplementary Appendix S2.

### A neural proxy for the model-based latent decision variable

Next, we leveraged human EEG recordings to establish a neural proxy for the model-based latent decision variable of the DDM. Previous research has shown that preparatory motor signals, such as the Lateralized Readiness Potential (LRP), encode this latent decision variable^25–31^. This means that we can use the LRP to validate the neural plausibility of the extended DDM.

In our EEG sample (N = 45), we observed a clear correspondence between the LRPs and latent decision variables of the DDM. Specifically, motor preparation demonstrated an immediate build-up towards the correct response on congruent trials, while exhibiting a “dip” towards the incorrect response (i.e., irrelevant capture) before favoring the correct response on incongruent trials (Fig. 3A). Next, we extracted features from these LRP waveforms (see Methods for details) to test whether they function as neural proxies for the DDM parameters. First, we extracted the average “LRP slope” as a proxy for the drift rate. Second, the positive peak amplitude of the LRPs, referred to as “LRP amplitude”, was extracted to approximate the decision boundary. Third, the onset latency of the LRPs (“LRP latency”) can function as a surrogate for the non-decision time. Last, the early negative peak amplitude on incongruent trials, denoted as “LRP dip” was chosen as a neural proxy for the degree of model-based irrelevant capture. As a first proof that these neural indices closely matched their model-based counterparts, we showed that these neural indices showed similar logarithmic (LRP slope: β = -0.139, *SE* = 0.031, *t*(44) = -4.48, *P* < .001; LRP amplitude: β = -0.107, *SE* = 0.029, *t*(44) = - 3.69, *P* < .001; LRP latency: β = -0.101, *SE* = 0.044, *t*(44) = -2.27, *P* = .029) or linear (LRP dip: β = -0.227, *SE* = 0.035, *t*(44) = -6.35, *P* < .001) decreases over the course of the experiment, mirroring their model-based counterparts (Fig. 3B).

**Fig 3.**
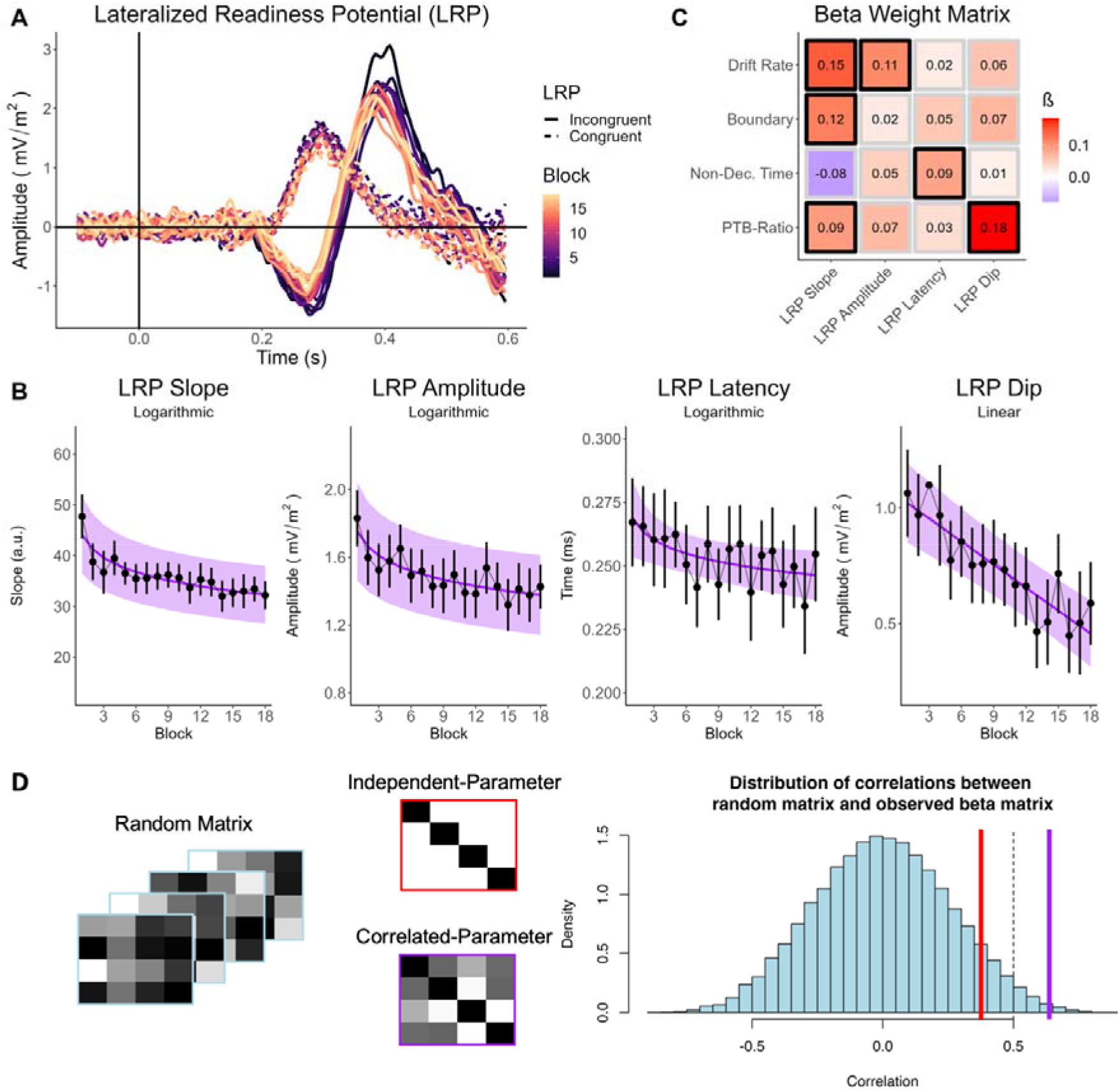
Preparatory motor signals and their relationship to the diffusion model. **(A)** Lateralized Readiness Potential (LRP) for congruent trials (dashed lines) and incongruent trials (solid lines) and each block. **(B)** Time courses of the extracted LRP features. The subtitle denotes the best model for time-on-task (linear, logarithmic, quadratic or cubic) based on BIC. **(C)** The (standardized) beta weights from the multivariate mixed effect model quantifying the intra-individual relationships between local fluctuations in LRP features and local fluctuations in DDM parameters. Black boxes denote coefficients where the 95 % HDI does not contain zero. **(D)** Examples of random model matrices used for the representational similarity analysis (blue outline), the independent-parameter model matrix (red outline), the correlated-parameter model matrix (purple outline) and the null distribution based on the random model matrices (blue) and the observed correlations for the specific (red line) and overlap (purple line) model. The dashed line refers to P = .05. PTB-Ratio = peak-to-bound ratio. Error bars represent the 95% within-subject CI.

While the similarity between the model parameters, neural indices and their underlying latent decision variables can already be appreciated qualitatively, we also sought to quantitatively evaluate this. To this end, we examined the intra-individual similarity between the time series of the DDM parameters and the neural indices. Specifically, we used a Bayesian Multivariate Linear Mixed Effects Model to predict local fluctuations (i.e., the detrended or residual timeseries, see Methods) of the DDM parameters by local fluctuations of the neural indices. This model returns a beta matrix, which should not be confused with a correlation matrix, as it quantifies the hierarchically estimated within-subject relationships with the effect of the other independent variables removed and the correlation across dependent variables taken into account. As expected, we found positive relationships between LRP slope and drift rate (Fig 3C), β = 0.145, 95% HDI (Highest Density Interval) [0.053, 0.231], *pd* (probability of direction, i.e., proportion of the posterior distribution that is of its median sign) > 99.99%, LRP latency and non-decision time, β = 0.085, 95% HDI [0.009, 0.159], *pd* = 98.77%, and LRP dip and peak-to-bound ratio, β = 0.183, 95% HDI [0.092, 0.268], *pd* > 99.99%, showing that fluctuations in model parameters closely tracked fluctuations in the neural indices. We did not find a relationship between LRP amplitude and boundary, β = 0.022, 95% HDI [-0.068, 0.115], *pd* = 67.79%. Yet, boundary was related to the LRP slope, β = 0.116, 95% HDI [0.017, 0.208], *pd* = 99.09%, as was peak-to-bound ratio, β = 0.092, 95% HDI [0.005, 0.178], *pd* = 98.28%.

While this beta matrix shows the relationships that are present between the DDM parameters and the neural indices, it does not necessarily show whether these results adequately portray the expected overlap between the two domains. Therefore, we adapted a model comparison approach using a representational similarity analysis logic. To evaluate whether the overlap is larger than what could be expected on the basis of chance, we constructed a null distribution by simulating fifty thousand random beta matrices and we computed the Spearman correlation between the random matrix and the observed beta matrix (Fig. 3D). Next, we created two theoretical models, which represent the degree of overlap one would expect between the DDM and neural domain. In a first model, the “independent-parameter model”, we assume that drift rate is specifically related to LRP Slope, decision boundary only to the LRP amplitude, non-decision time only to the LRP latency and PTB-Ratio only to the LRP Dip. This model can thus be represented as the diagonal (Fig. 3D). However, we know that this assumption of independence is not realistic because of the existing correlations between the DDM parameters themselves, which can be expected to manifest in the neural data as well (e.g., with an established correlation between drift rate and boundary, one also expects a correlation between LRP slope and LRP amplitude). Therefore, in a second model, the “correlated-parameter model”, we included the empirically observed correlations between the DDM parameters in the representation model (Fig. 3D). Finally, we evaluated the representational similarity between the observed beta matrix and our two model matrices through Spearman correlation. Our findings indicate that while the correspondence between the observed data and the independent-parameter model did not demonstrate a significant improvement over the null (random) model (*R* = .376, *P* = .076), the correlated-parameter model significantly outperformed the null model (*R* = .637, *P* = .005).

Together, this indicates that considering the observed overlap between DDM parameters, there is a relatively strong convergence between neural indices and DDM parameters. This represents a novel and crucial step in establishing the neural plausibility of this DDM for the Flanker task^24^. Analyses on the inter-individual level can be found in Supplementary Appendix S3.

### Metacognitive experience and model-based task performance

Having established the computational basis of task performance, validated by our neural proxy, we now aimed to answer our main question: which subjective experiences provide metacognitive insight in which objective model-based task performance parameters? First, we established the time-on-task effects for the metacognitive experiences. The average level of conflict aversiveness was found to be significantly above zero, *b_0_* (intercept) = 16.52, *SE* = 2.11, *t*(110) = -7.84, *P* < .001, indicating that participants generally found conflict aversive, consistent with previous findings^12^. Although the best time-on-task model for conflict aversiveness was logarithmic (Fig. 4A), there was no significant change in conflict aversiveness with time-on-task, β = -0.008, *SE* = 0.028, *t*(110) = -0.20, *P* = .839. However, a more detailed exploration of inter-individual differences, as presented in Supplementary Appendix S4, reveals a more nuanced picture. These analyses revealed the presence of two subgroups: one showing an increase in conflict aversiveness over time and another showing a decrease. Importantly, the effects within these subgroups offset each other when analyzed at the group level, leading to the appearance of no significant change. Boredom increased in a quadratic fashion, β*_linear_* = 0.300, *SE* = 0.025, *t*(110) = 11.82, *P* < .001, β*_quadratic_* = -0.059, *SE* = 0.010, *t*(1775) = --5.89, *P* < .001, showing an initial steady increase that slowed down towards the end of the experiment. Effort showed a small, but constant linear increase, β = 0.128, *SE* = 0.029, *t*(110) = 4.39, *P* < .001. Fatigue showed a cubic pattern: the initial increase in the beginning of the experiment slowed down in the middle of the experiment but then increased again more steeply towards the end of the experiment, β*_linear_* = 0.361, *SE* = 0.035, *t*(264.20) = 10.27, *P* < .001, β*_quadratic_* = -0.074, *SE* = 0.009, *t*(1774) = -8.21, *P* < .001, β*_cubic_* = 0.118, *SE* = 0.023, *t*(1774) = 6.90, *P* < .001. Frustration showed a quadratic pattern, β*_linear_* = 0.184, *SE* = 0.027, *t*(110) = 13.88, *P* < .001, β*_quadratic_* = -0.056, *SE* = 0.009, *t*(1775) = -6.22, *P* < .001, with an initial steady increase that plateaued and eventually even seemed to reverse towards the end of the experiment. Together, these time-on-task effects show that the different metacognitive experiences had distinct temporal dynamics.

**Fig 4.**
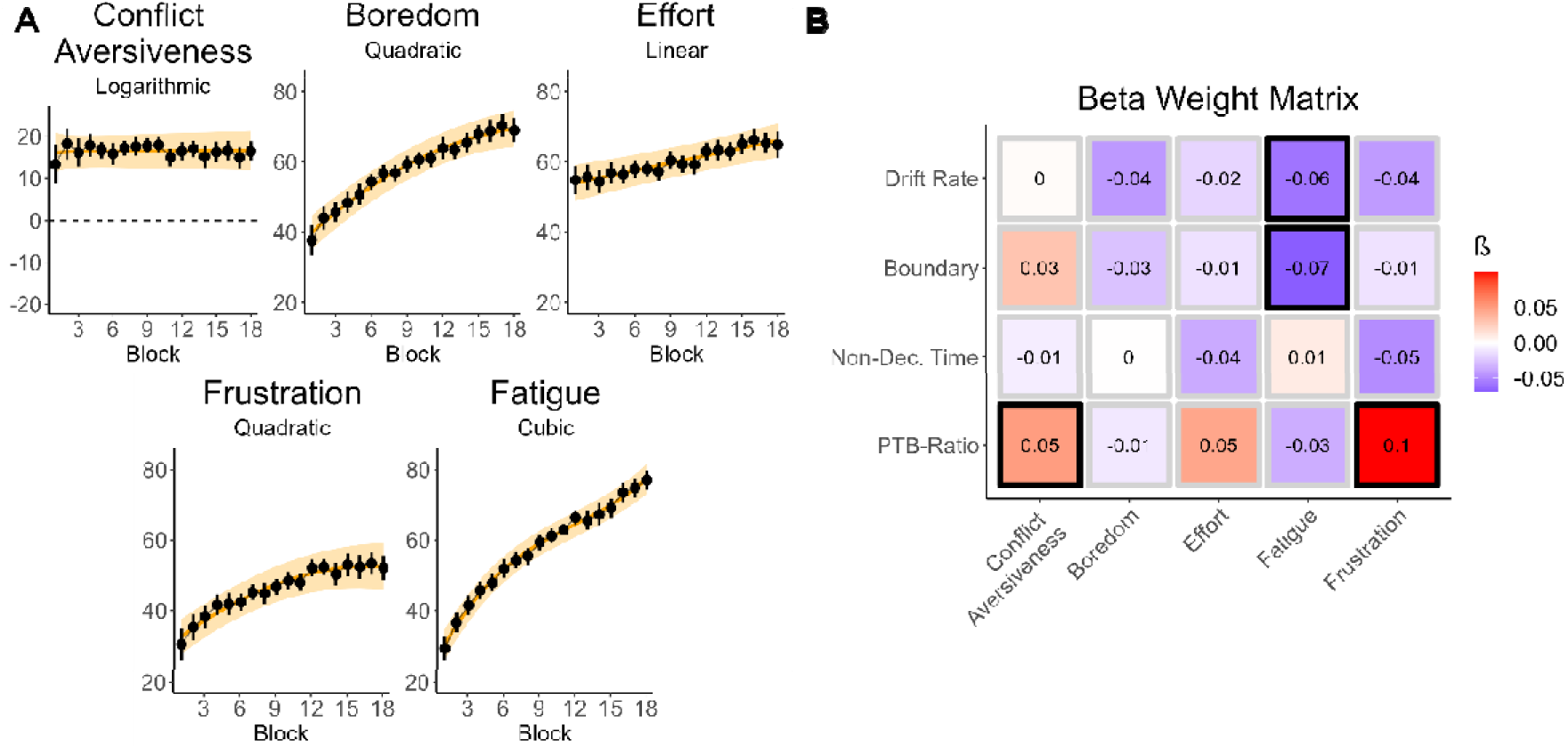
**(A)** Time-on-task effects for the metacognitive experiences. **(B)** Intra-individual relationships between the DDM parameters and Metacognitive Experiences. Black boxes denote coefficients where the 95 % HDI does not contain zero. The subtitle denotes the best model for time-on-task (linear, logarithmic, quadratic or cubic) based on BIC. PTB-Ratio = peak-to-bound ratio. Error bars represent the 95% within-subject CI.

Next, we quantified the relationships between metacognitive experiences and the DDM parameters, within individuals, by conducting a Bayesian Multivariate Mixed Effects Model predicting local fluctuations in DDM parameters by local fluctuations in metacognitive experiences (for the inter-individual level analyses see Supplementary Appendix S4, for the intra-individual level analyses between the behavioral indices and metacognitive experiences see Supplementary Appendix S5). Our analysis yielded two primary findings. First, peak-to-bound ratio was positively related to conflict aversiveness, β = 0.050, 95% HDI [0.004, 0.092], *pd* = 98.54%, and frustration, β = 0.099, 95% HDI [0.048, 0.149], *pd* > 99.99%, showing that people evaluated conflict as less negative and were less frustrated when the irrelevant stimulus dimension captured their attention less (Fig. 4B). Second, drift rate displayed a negative relationship with fatigue, β = -0.059, 95% HDI [-0.108, -0.011], *pd* = 99.23%, as did boundary, β = -0.069, 95% HDI [-0.121, -0.018], *pd* = 99.58%: higher levels of fatigue were associated with less efficient evidence accumulation and lower levels of cautiousness, indicating more general changes in performance. Together, our findings show that frustration and conflict aversiveness offer metacognitive insights specifically into the extent of irrelevant capture, while fatigue provides a more general insight into performance efficiency, along with one’s level of response caution.

### A rational perspective on task performance

To unravel the underlying mechanisms responsible for the observed dynamics of task performance we turned to a rational framework. The strongest time-on-task effect in performance was found on the decision boundary, which strongly decreased over time. Notably, within sequential sampling frameworks, the decision boundary stands out as a distinctive parameter intimately linked to strategic control, whereas the other parameters are predominantly thought to reflect ability^32^ (but see^33^). Crucially, setting the decision boundary, and thus strategic adjustments in task performance, directly impacts the optimality of decision making^34–37^. Specifically, there usually is a single setting for decision boundary that will maximize an objective criterion such as the reward rate^34^. Setting the decision boundary too low results in faster sampling of rewards but at a lower success rate, while setting it too high will lead to more successes at the cost of time. The optimal setting of the decision boundary, therefore, aims to maximize reward in the least amount of time.

However, in the current conflict task and model, the maximization of an objective criterion such as the (pseudo-)reward rate (in the absence of rewards, as in our task, correct responses can be considered pseudo-rewards^38^) will also necessarily depend on the strength of the irrelevant dimension, as modeled by the peak amplitude parameter (which showed the second largest time-on-task effect). This dependence of optimality on irrelevant capture is demonstrated by the simulations performed in Fig. 5A. This shows that in order to optimize the reward rate, one needs to find the setting of the decision boundary that is sufficiently high, relative to the strength of the irrelevant dimension. Consequently, when the sensitivity to irrelevant capture decreases over time, as we observed, it can be considered rational to lower the decision boundary to a similar degree. Failing to adapt the decision boundary as irrelevant capture decreases would result in unnecessary time spent gathering rewards (i.e., correct responses). In line with this, we found a very strong positive relationship between local fluctuations in peak amplitude and local fluctuations in boundary, β = 0.829, 95% HDI [0.796, 0.862], *pd* = 100%, showing that changes in peak amplitude were always accompanied by highly similar changes in boundary.

**Fig 5.**
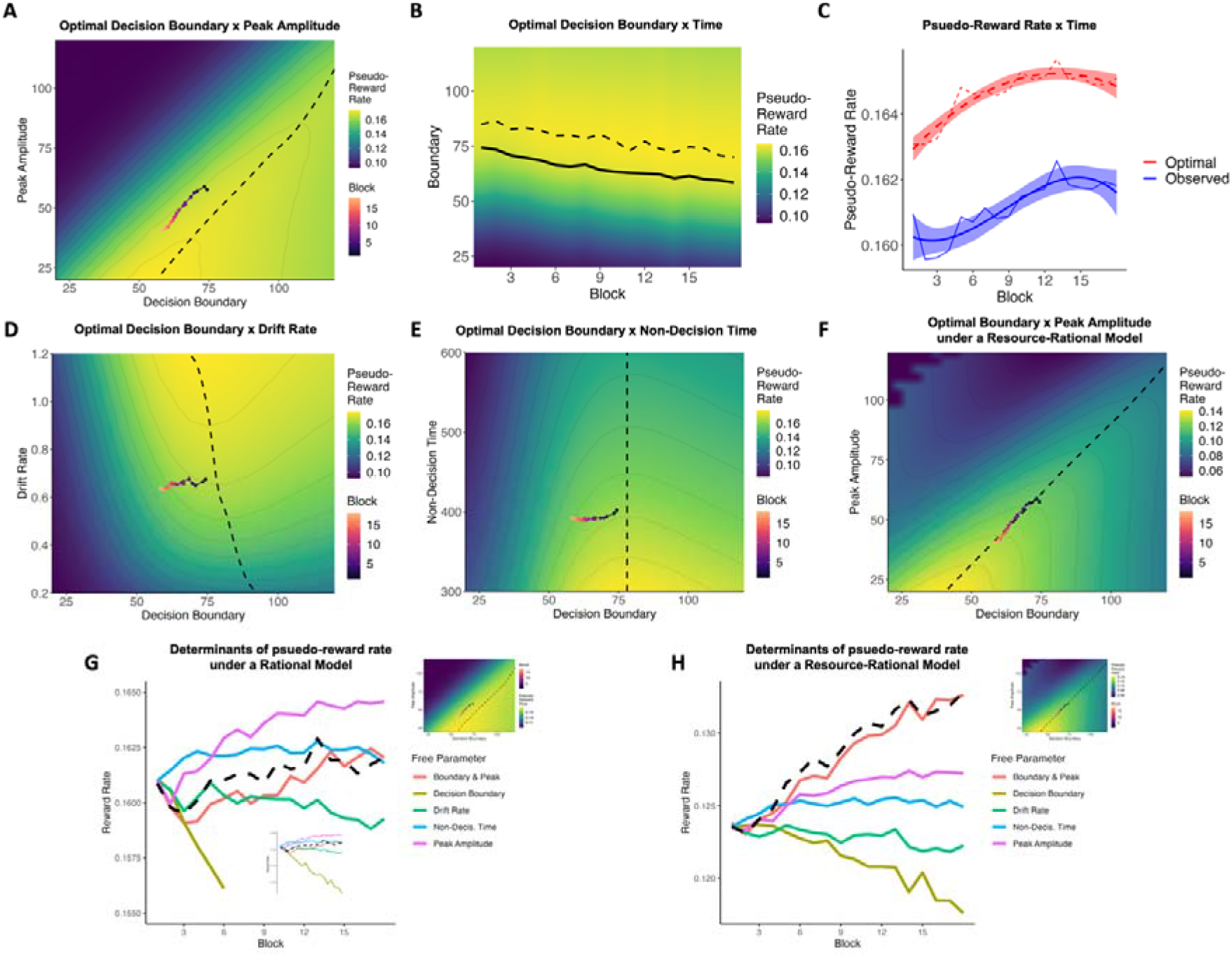
**(A)** Pseudo-reward rate landscape for peak amplitude and boundary, revealing the optimal setting of the decision boundary for each level of peak amplitude (dashed line). The colored line depicts the observed time-on-task trajectory of peak amplitude and boundary within this space. **(B)** The pseudo-reward rate landscape for boundary across blocks – given the trajectory of the other parameters. The dashed line shows the optimal boundary trajectory, while the solid line shows the observed boundary trajectory. **(C)** Optimal versus observed changes in pseudo-reward rate with time-on-task. **(D)** Pseudo-reward rate landscape for drift rate and boundary (E) and non-decision time and boundary. **(F)** Pseudo-reward rate landscape for peak amplitude and boundary within a resource-rational framework. **(G)** Determinants of reward rate in the rational model. Each line shows the trajectory of pseudo-reward rate if only the free parameter was allowed to vary over the experiment while the others are fixed to their initial value. **(H)** Determinants of pseudo-reward rate in the resource-rational model. Error bars represent the 95% within-subject CI.

Further, when simulating the optimal decision boundary based on the observed trajectory of the other parameters, we found that lowering the decision boundary over time (Fig. 5B, dashed line) was indeed optimal, and participants appeared to follow a similar pattern. Similarly, the optimal and observed trajectory in the pseudo-reward rate landscape of boundary and peak amplitude (Fig. 5A) further confirmed that similar decreases in peak amplitude should be and are followed by similar decreases in boundary. Finally, we show that lowering the decision boundary was indeed beneficial, as it led to an increase in the pseudo-reward rate with time-on-task (Fig. 5C) for both the optimal trajectory (best model: logarithmic), β = 0.903, *SE* = 0.107, *t*(16) = 8.41, *P* < .001, and the observed trajectory (best model: cubic), β*_linear_* = 0.818, *SE* = 0.112, *t*(14) = 7.31, *P* < .001; β*_quadratic_* = -0.222, *SE* = 0.112, *t*(14) = -1.98, *P* = .067; β*_cubic_* = -0.326, *SE* = 0.112, *t*(14) = -2.91, *P* = .011.

To test whether changes in peak amplitude inform adjustments in decision boundary, we conducted a Granger causality analysis to examine temporal flow of information between these variables. We found that past values of peak amplitude from the previous block predicted decision boundary above and beyond past values of boundary (z[ = 3.53, P < .001) but not the other way around (z[ = 1.08, P = .281). It is important to note that while Granger causality analysis provides insights into the temporal associations between variables, it does not establish causal relationships definitively. We verified that the relationship between peak amplitude and boundary was not a modeling artifact, as it did not arise during parameter recovery (R = -.40, P = .700). We also ruled out alternative models of rationality. We show that the dynamics of decision boundary can only be considered rational with respect to the dynamics of peak amplitude (Fig. 5A). With respect to drift rate and non-decision time, the dynamics of the boundary do not follow the optimal trajectory, even exhibiting orthogonality (Figure 5D-E).

We further show that the observed increase in pseudo-reward rate primarily arises from the interplay between peak amplitude and decision boundary (Fig. 5G). By simulating various pseudo-reward rate trajectories, each time isolating the effect of an individual parameter while keeping the others fixed to their initial value, we found that the decrease in peak amplitude significantly boosts the reward rate, while the observed changes in drift rate and non-decision time had smaller effects. Notably, the reduction in decision boundary reduces the reward rate, underscoring its rationality only when viewed in the context of peak amplitude. When viewed together, they closely align with the observed reward rate trajectory.

Interestingly, these simulations also reveal the potential suboptimality of our subjects, as they consistently fall below the optimal solution (Fig. 5C). This behavior could be interpreted as a form of resource-rational behavior, where participants demonstrate rationality to a certain extent, avoiding excessively high decision boundaries that only marginally increase the reward rate. In essence, there seems to be an additional cost currently not reflected in the pseudo-reward rate. One way to address this behavior, is to introduce an additional cost to the model which we hypothesized to be equivalent to the height of the decision boundary (scaled by a weight). This effectively creates a resource-rational model where the unnecessary slowing down associated with high boundaries is avoided (cf. participants behavior). Integrating this cost in the rational models revealed its ability to account for the perceived suboptimality (Fig. 5F). Moreover, this model again highlights the collaborative impact of peak amplitude and decision boundary on pseudo-reward rate. Assuming perfect optimality among participants within the resource-rational model, the observed dynamics of peak amplitude and decision boundary work together to have the strongest influence on the increase in pseudo-reward rate (Fig. 5H).

Together, these optimality analyses show that the main temporal dynamics of task performance can be understood as two sides of the same coin: an increased ability to filter out irrelevant information (decrease in Peak Amplitude) supports strategic adaptations (lowering the Boundary) to maximize the pseudo-reward rate.

### Metacognitive experiences track distinct rational adjustments in task performance

In a final set of analyses, we aimed to bring together how the various metacognitive experiences could be related to the main performance optimization dynamics identified in the previous section. To investigate this question, we conducted multilevel path analyses using Bayesian Multivariate Linear Mixed Effects Modeling where we tested whether the relationship between peak amplitude and boundary is mediated by the metacognitive experiences. It is important to clarify that this path model does not represent a definitive causal account of the current findings but is meant to provide an exploratory narrative framework that integrates our previous findings and makes specific predictions that could be tested in future studies.

In a first path model, we performed parallel mediation to test whether any of the metacognitive experiences mediated the relationship between peak amplitude and boundary. Here, we found that peak amplitude predicted conflict aversiveness, β*_direct_* = 0.052, 95% HDI [0.007, 0.096], *pd* = 100%, while effort and frustration predicted the decision boundary (β*_direct_* = -0.030, 95% HDI [-0.058, -0.001], *pd* = 97.7%; β*_direct_* = -0.052, 95% HDI [-0.082, -0.021], *pd* = 100%). However, we did not find evidence for a mediator. Given the unexpected outcome that fatigue did not predict boundary (in contrast to our prior findings), we constructed a second model to investigate whether boundary predicted fatigue instead. Our analysis revealed that boundary did indeed predict fatigue, β*_direct_* = -0.082, 95% HDI [-0.132, - 0.031], *pd* = 99.9%. In a third model, we introduced serial mediation to investigate whether peak amplitude’s prediction of conflict aversiveness, along with effort and frustration’s prediction of boundary, created an indirect pathway through these variables. Although we did not find evidence for an indirect path, we did identify a relationship between conflict aversiveness and effort, β*_direct_* = 0.064, 95% HDI [0.011, 0.115], *pd* = 99.2%. This led us to explore a fourth model, incorporating serial mediation from conflict aversiveness through effort and frustration. In this model, we discovered a significant indirect effect over these three metacognitive experiences, β*_indirect_* < -.001, 95% HDI [<-.001, <-.001], *pd* = 97.97%. In a fifth and final model, we examined whether there was a relationship between frustration and fatigue and whether this relationship was mediated by the decision boundary. We observed a direct relationship between frustration and fatigue, β*_direct_* = 0.257, 95% HDI [0.189, 0.326], *pd* = 100%, and we found that boundary mediates this relationship, β*_indirect_* = 0.003, 95% HDI [0.001, 0.008], *pd* = 99.70%. Leave-one-out cross-validation revealed that the final model (Model 5) outperformed the other models in terms of expected log pointwise predictive density (ELPD), which serves as a measure of predictive accuracy or fit. The differences in ELPD (and SE of the differences) between Model 5 and the other models were as follows: Model 1: -142.3 (22.4), Model 2: -174.9 (24.9), Model 3: -136.1 (19.8), Model 4: -108.0 (16.0). A visual representation of the final path model is depicted in Fig. 6 and the other path models can be found in Supplementary Appendix S6.

**Fig 6.**
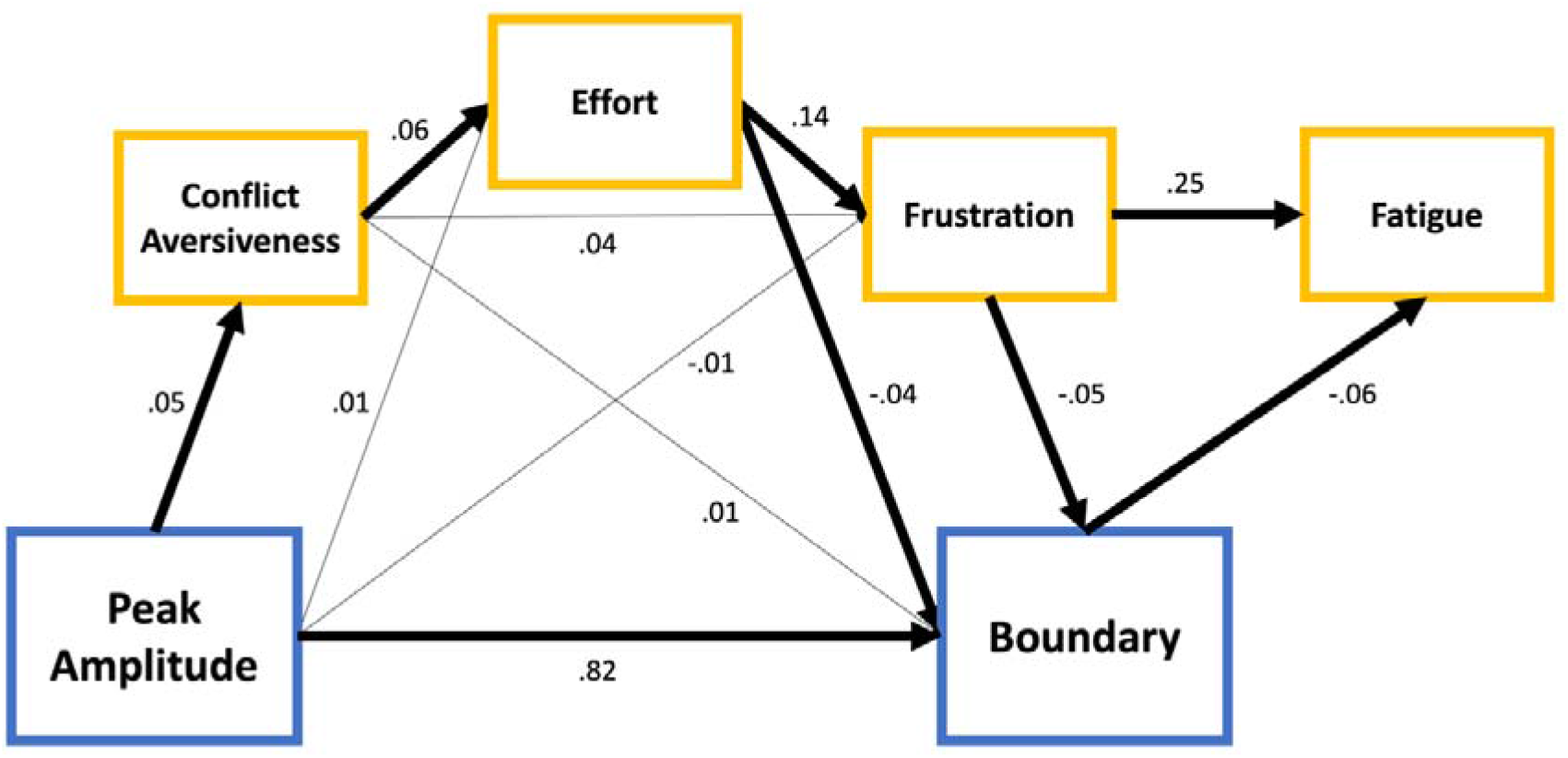
Path Model. Significant relationships (for which the 95% CI does not include zero) have dark arrows. Blue relates to task performance parameters, yellow to metacognitive experiences.

In summary, these findings suggest potential pathways through which various metacognitive experiences may track and influence rational adaptations in task performance. Conflict aversiveness appears to reflect a subjective assessment of irrelevant capture. This subjective assessment might then affect the level of effort exerted in the task. Local changes in effort could subsequently guide adjustments in the decision boundary, either directly or through feelings of frustration. Furthermore, the ongoing effort required to adapt the decision boundary may contribute to feelings of fatigue.

## Discussion

The current work investigated how metacognitive experiences dynamically track cognitive processes underlying task performance. To this end, we employed a time-on-task design during which participants performed a conflict Flanker task for two hours where irrelevant information had to be ignored. This task was structured in short blocks, and after each block, we probed participants’ metacognitive experiences of conflict aversiveness, boredom, effort, fatigue and frustration.

We showed that the temporal dynamics of task performance were dominated by two shifts: a reduction in irrelevant capture and a decrease in decision boundary as time-on-task progressed. Next, we demonstrated that changes in the model parameters over time closely resembled changes in the LRP indices over time, thereby confirming the biological plausibility of our model. Subsequently, we found that these main time-on-task dynamics in performance were strongly related to each other and appeared to reflect two sides of the same coin: a rational attempt to optimize the (pseudo-)reward rate. Metacognitive experiences closely tracked these adaptive shifts. Conflict aversiveness mirrored changes in irrelevant capture, thereby reflecting shifts in the potential reward rate, while frustration and fatigue seemed to serve as input and output of a regulatory process of decision boundary adjustments, aimed to harness the potential reward rate but limited by the exerted effort. Collectively, these findings highlight the link between metacognitive experiences and the regulation of cognitive processes, proposing potential pathways through which these experiences could impact adaptive behavior by monitoring and possibly shaping the optimality of decision-making.

We found that conflict aversiveness closely tracks the objective level of interference due to conflict or irrelevant capture both at the intra- and inter-individual level (see Supplementary Appendix S4): higher levels of conflict aversiveness were associated with higher levels of model- and neural-based irrelevant capture and vice versa. This finding is consistent with the notion that the aversive experience of conflict mirrors the cost of exerting control, and these costs are proportional to the intensity of required control. In other words, higher control intensities incur greater costs^9–12,15,16^. This is in line with previous empirical investigation into the subjective nature of control processes demonstrating relationships between objective performance costs due to conflict, subjective estimates of this cost and subsequent avoidance behavior^39–42^. On the other hand, some studies have failed to establish such connections (e.g.,^43,44^), potentially because they examined interference effects in RT and error rates separately, without employing a comprehensive model-based estimate of interference that accounts for the positioning of one’s decision boundary (see below for more on the importance of incorporating measures of decision boundary for the computation of irrelevant capture).

Moreover, combining RT and accuracy naturally lends itself to a normative interpretation rooted in reinforcement learning and neuroeconomic frameworks^15,45^. That is, the degree of irrelevant capture can be conceptualized as the loss in reward rate due to conflict. In essence, when irrelevant capture is substantial, it imposes a significant cost on the potential reward rate achievable, as incongruent trials carry both a time penalty and an increased risk of failing to attain the (pseudo-)reward. This perspective aligns with reinforcement learning theories, which have posited that the aversive evaluation of conflict can be inferred from task performance based on the lowered reward expectancy for incongruent compared to congruent trials^46,47^. Furthermore, our finding that individuals started to perceive conflict more positive as irrelevant capture diminished over time aligns with the idea that successfully resolving conflict can be experienced as rewarding, or at least more positive, and this seems to be driven by changes in the underlying reward expectancies^47–54^.

It is essential to underscore the importance of having an accurate subjective estimate of the degree of irrelevant capture as this information plays a crucial role in guiding further behavioral adjustments. In fact, some have argued that these subjective estimates may hold even greater importance in driving adaptations than the objective level of interference alone^9,55,56^. Particularly, when levels of irrelevant capture shift over time, individuals must adapt their decision boundary in response to these changes to maintain optimal performance, as demonstrated by our simulations. However, our results do not show that such subjective estimates (e.g., conflict aversiveness) necessarily promote strategic adaptations (aligning with mixed findings regarding the role of subjective estimates in guiding trial-by-trial adaptations^57–59^). Instead, effort and frustration seem to mediate the strategic changes, and these changes were associated with feelings of fatigue.

The finding that the experiences of effort, fatigue and frustration are related to local changes in decision boundary are in line with prior research^14,60^. For example, Lin et al.^14^, found that an ego depletion manipulation resulted in lower decision boundaries, which were linked to more pronounced feelings of fatigue and frustration. Even more, their study showed that higher perceived effort predicted greater reductions in the decision boundary. Similarly, in our study, we observed comparable patterns: over time, global trends showed that participants consistently reported heightened levels of effort, frustration and fatigue, indicating they found the task demanding despite experiencing less irrelevant capture and improving performance as they implemented normative resource-rational adaptations (i.e., lowered their decision boundary and increased pseudo-reward rate). In other words, even successfully adapting behavior to optimize performance can be experienced as fatiguing when it requires significant effort. Towards the end of the experiment, participants did begin to exhibit performance decrements, as evidenced by a decrease in pseudo-reward rate, suggesting they were approaching their cognitive capacity limits. While our study replicates the relationships reported by Lin et al., we argue against expecting this relationship to be consistently in the same direction across different tasks or contexts. We propose that any adjustment, whether it involves becoming “more reckless” (lowering the decision boundary) or “more cautious” (raising the decision boundary), could be related to feelings of fatigue and frustration, depending on the normative context.

The reason for this claim is that in our view, fatigue arises as a consequence of the effortful process of adapting behavior in an optimal manner. Although participants appear successful at dealing with the task at hand (they were able to increase the pseudo-reward rate), it is unclear whether this made the task easier for them. Despite improved performance, participants consistently reported heightened effort levels, suggesting that their enhanced performance was tied to an ongoing, effortful process of adaptation. Even more, towards the end of the experiment, participants began to exhibit performance decrements, indicated by a decrease in pseudo-reward rate, suggesting they were approaching their capacity limit. Together, these observations underscore the demanding nature of the task and the effort required for sustained performance optimization.

The normative models also revealed the suboptimality of our participants. Intriguingly, in contrast to the finding that humans often adopt strategies that are overly cautious (with decision boundary above the optimal boundary, e.g.,^34,61,62^), our simulations reveal that participants leaned towards an overly urgent strategy (with decision boundary below the optimal boundary^63^). This suggests that participants were more inclined to reduce the cost of time than maximize the absolute reward rate. This finding aligns with resource-rational frameworks^16,64–69^. It appears that the perceived cost of time associated with allocating control resources (which we could successfully model as a cost of the boundary) dissuades individuals from investing the additional resources necessary to achieve the maximum reward rate, leading to a tradeoff between effort and accuracy. This tradeoff phenomenon has been observed beyond our study in various cognitive domains such as visual cognition^70^ and language^71^.

Our investigation also uncovered a previously overlooked aspect concerning the measurement of conflict. Our findings imply that the extent of irrelevant capture not only depends on the strength of evidence from the irrelevant dimension but also on the height of one’s decision boundary. Thus, researchers may have inadvertently mixed measures of irrelevant capture with measures of cautiousness, potentially explaining the low correlations of irrelevant capture measures in individual difference and reliability studies across various conflict tasks^72–74^. An unconfounded measure of irrelevant capture should consider both the impact of the irrelevant dimension and express this impact relative to the height of one’s decision boundary. For instance, if a researcher aims to correlate peak amplitude parameter values from multiple conflict tasks to assess a general sensitivity to irrelevant information, disregarding the height of the decision boundary can lead to misleading results, as the decision boundary is likely to vary between tasks, affecting the true impact of the peak amplitude parameter on decision formation.

Further, we suggest a temporal dimension to the relationship between peak amplitude and decision boundary in the current experiment: changes in peak amplitude preceded adjustments in decision boundary. While our data do not allow for definitive causal claims, we believe most empirical evidence points in one direction. Moreover, there are reasons to assume that it is more plausible that people adapt their decision boundary rather than their peak amplitude. The decision boundary is typically considered a strategic parameter that can be manipulated via instructions^81^ and in response to conflict^33,82–87^. On the other hand, behavioral measures for irrelevant capture (interference effects, which are here quantified by peak amplitude) are often utilized to measure individual differences in inhibitory capacity or executive functioning, implying a closer association with one’s ability rather than strategic adjustments^74,88–91^. Second, within a sequential sampling framework, the concept of optimality is typically applicable solely to the decision boundary^34^. When evaluating an objective criterion like reward rate, an optimal decision boundary can be identified with respect to the other parameters, whereas there isn’t an optimal drift rate, peak amplitude, or non-decision time with respect to the other parameters. Higher drift rate and lower peak amplitude and non-decision time are always more advantageous.

To conclude, our study revealed multiple dynamic interactions between several metacognitive experiences and cognitive processes during prolonged task engagement. By combining cognitive modeling with neural measures, we provide novel and insightful perspectives into the intricate interplay between metacognition and cognition by offering specific pathways through which metacognitive experiences might interact with task performance. These findings have implications across various domains such as education, clinical contexts, and professional challenges. For instance, tailored interventions aimed at cultivating frustration awareness while reducing effort awareness, could enhance decision-making processes in challenging situations. Our study also offers valuable insights for normative computational models of subjective metacognitive experiences^17,18,75–80^.

## Methods

### Participants

Seventy-one participants participated in the online behavioral experiment which was conducted on Prolific Academic and took on average 117.7 minutes to complete (participants were compensated at 15.12 pounds in total or 7.56 per hour). All participants performed the experiment in English (only participants whose first language was English were allowed to participate). Five participants were excluded from the analyses due to not adhering to the instructed break time (i.e., 30 seconds, more than 2.5 SD above the mean break time), missing too many responses (more than 2.5 SD above the mean number of missed responses), responding too slow (higher than 2.5 SD above the mean RT) or making too many errors (higher than 2.5 SD above the mean error rate). The average age of the remaining 66 participants (31 males, 33 females, 2 unknown) was 29.73 years (SD = 5.26, min = 18, max = 36). Participants had to agree with a consent form before starting the experiment. Forty-nine participants participated in the EEG experiment which was conducted at the Research Center for Mind, Brain and Behavior at the University of Granada and took on average 116.8 minutes to complete (participants were compensated 35 EUR for the whole session, including preparation time). Four participants were excluded from the analyses due to not adhering to the exclusion criteria specified above. The average age of the remaining 45 participants (16 males, 29 females) was 21.42 years (SD = 2.75, min = 18, max = 31). Thirty-nine participants performed the experiment in Spanish, and 6 participants performed the experiment in English. Participants signed an informed consent form before starting the experiment. All experiments were approved by the Ethical Committee of Ghent University Psychology and Educational Sciences or University of Granada.

### Design, Paradigm and Procedure

The experiment was programmed using JsPsych^92^ in the online setting and PsychoPy 3 ^93^ in the EEG setting. Participants were instructed that they were going to perform a simple decision-making task almost continuously over a period of two hours. In the online setting, they were encouraged to turn off notifications from their smartphone and computer. Participants were instructed that they should not miss more than 5 % of the trials, make more than 30 % errors or respond too fast (response < 100 ms) on more than 5 % of the trials.

Next, participants were instructed on the Flanker task (based on^25^) which stated that they had to press S (online) or left control (EEG) when the central target arrow pointed to the left, and press L (online) or right control (EEG) when the central target arrow pointed to the right. Participants started each trial with a fixation display (mean = 700 ms, randomly drawn from a uniform distribution between 600 and 800 ms). Next, the horizontal flankers appeared for a short duration (83 ms) followed by the central target (33 ms). After this, the stimulus disappeared from the screen and participants were required to respond within a 1200 ms window (see Fig. 1 for a visual depiction of the timing and sequence of events in the Flanker task). They were instructed to respond as fast and accurately as possible. The next trial started immediately after a response was made. This was followed by a short practice version of the Flanker task (60 trials) where trial-by-trial feedback was provided (“Too slow”, “Too fast”, “Correct!”, “Wrong!”) ending in a display of feedback showing their average RT and accuracy, the percentage of missed responses and the percentage of impossibly fast responses. Participants needed to achieve an accuracy of 70 % in this practice phase to continue the experiment.

Next, participants were told that they would be questioned about their mental state after each block of the Flanker task. Participants were told that their feelings towards the task and stimuli might change over the course of the experiment, but they do not necessarily have to and that it is simply important to provide an accurate and reliable report of their subjective experience at each point in time. Next, participants were explained the difference between congruent and incongruent stimuli, so they could give their informed preference about whether they preferred being presented with congruent (“same”) or incongruent (“different”) stimuli (i.e., conflict aversiveness). A Visual Analogue Scale (VAS) was presented that started in the center (which was labeled as indifferent) and had “extreme preference for SAME” and “extreme preference for DIFFERENT” on each end of the VAS (left and right respectively, see Fig. 1 for a simplified visual depiction). Next, they received VAS polling their levels of effort (“How hard did you work in the task?”; not hard – very hard), frustration (“How annoyed are you with the task?”; not annoyed – very annoyed), boredom (“How boring do you find task?”; not boring – very boring) and fatigue (“How mentally fatigued are you now?”; not fatigued – very fatigued). These questions were based on previous work^14^.

Finally, participants started the actual experiment, which consisted of 18 blocks of 260 trials each in which congruency (congruent, incongruent) and target direction (left, right) were always balanced. After each block, participants received VAS polling their metacognitive experience and participants were allowed to take a short break of 30 seconds (and were instructed to strictly adhere to this break time).

### Time-on-task Analyses

All behavioral and modeling analyses were performed in R (4.1.3). For the raw RT analyses, we removed the first trial of each block and errors. We also removed trials with a RT below 200 ms (note that RT measurement was locked to the Flanker onset, which preceded target onset) and only kept trials between the 1^st^ and 99^th^ quantile (this rather minimal data trimming procedure was applied because of the subsequent diffusion modeling which models the whole RT distribution). For the metacognitive experience measures, we recoded the conflict aversiveness measure to increase interpretability. The responses to this scale were originally recorded from 0 to 100, with 50 being the indifference point. We rescaled this to vary from -50 to 50 so that zero became the indifference point, and positive values align with a more negative evaluation of conflict (a preference for congruent trials) and negative values align with a less negative evaluation of conflict (a preference for incongruent trials). In this way, higher values of conflict aversiveness denote a stronger disliking of incongruent stimuli. All other VAS were recorded from 0 to 100.

The behavioral data was further modeled using the Diffusion Model for Conflict Tasks (^24^) using the DMCfun package^94^. The model was fit on each block of data (260 trials) using the Root Mean Square Error cost function that quantifies the discrepancy between the model predictions and observed data on 19 bins (percentiles) of the correct RT distributions for congruent and incongruent trials and 5 bins of the Conditional Accuracy Function (CAF, see ^24^ and ^94^ for more details on the cost function). As there is no analytic solution for this diffusion model, predictions of the model were retrieved using Monte Carlo simulation and were based on 20000 trials for each congruency condition. The cost function was minimized using the Differential Evolution algorithm, which was ran for 500 iterations (fitting one dataset, i.e., one block of one subject, took roughly 45 minutes, resulting in a total of 61 days of serial computing time). We used the following boundaries for the parameter space: drift rate (0.2 – 1.2), decision boundary (30 - 110), non-decision time (200 – 600), peak amplitude (10 – 110). Peak latency was fixed to 28 and variability in non-decision time was fixed to 25. These fixed values are based on grand average values that were received after fitting the model per three blocks rather than on every block. We fixed these parameters as they were not our main interest (peak latency is mostly related to differences between conflict tasks, not within a given task) and lead to lower parameter recovery performance when fitting the model per block (rather than per three blocks, see Supplementary Appendix S1 for parameter recovery and fit assessment). The diffusion constant was fixed to 4 (cf.^24^). Note, that the plausible parameter ranges for this model differ from the standard DDM. Peak-to-bound ratio (PTB-ratio) was calculated by dividing the peak amplitude by the decision boundary.

To investigate the effect of Time-on-task (TOT), we ran Linear Mixed Effects model predicting the DDM parameters, the metacognitive experiences and the EEG indices (see below) by the effect of time-on-task (i.e., block: 1-18). We included random intercepts and random slopes for the effect of time-on-task up to the first degree (either linear or logarithmic depending on the best model). We tested multiple models of time-on-task (linear, logarithmic, quadratic and cubic) and chose the best model based on the lowest BIC. All performed statistical tests are two-sided.

### Intra-Individual Relationships

To investigate the intra-individual relationships between the time series of the DDM parameters and the metacognitive measures we used Multivariate Bayesian Linear Mixed Effects models (using the brms package^95^, MCMC settings: 8 chains of 4000 iterations with 1000 as warmup) to predict all the DDM parameters’ time series by all the metacognitive experience’ time series while controlling for Experiment (Behavioral, EEG). We included random slopes for all predictors but did not need random intercepts, as the mean of all time series were constrained to be zero due to standardization of each subjects’ time series. This allowed us to exclude inter-individual difference from this analysis (see Supplementary Appendix S3 and S4) and conveniently returns standardized beta weights which allows for a comparison of the relative magnitude of the different predictors. We also modeled the correlational structure between the dependent variables by including estimations of the correlations between the random slopes of the different dependent variables as well as correlations between the residuals of the multiple dependent variables. Weakly informative priors were used for all estimated beta weights (normal distribution with mean zero and a standard deviation of 2.5). We report the mean of the posterior distribution as the estimate for the standardized beta weight and its 95% Highest Density Interval (HDI). In addition, we report the probability of direction (pd), which is considered an index of effect existence and represents the certainty with which an effect goes in a particular direction ^96^. A *pd* value of 97.5% corresponds approximately to the two-sided *p*-value of .05.

These models were conducted on the detrended timeseries, or what we coin “local fluctuations”. The observed data was detrended by subtracting the best linear or logarithmic trend (depending on the best fit, as discussed above) from the observed data, i.e., leaving us with the residual time series. Thus, the beta weights reflect similarities in terms of sub-trend fluctuations between two given time courses – showing relationships that cannot be distorted by similarities in the global trend.

### Optimality Simulations & Path Analyses

The reward rate landscapes for the optimality analyses were performed by calculating the pseudo-reward rate for different combinations of peak amplitude (20-120, in increasing steps of .1) and decision boundary (20-120) given the average values for the other DDM parameters (Fig. 4A) or a range of decision boundary (20-120) for each block (Fig. 4B) given the other DDM parameters for that block (Fig. 4B). The pseudo-reward rate was calculated in the following manner (cf.^34,97^):

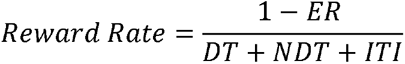

Where ER refers to the error rate, DT to the decision time, NDT to the non-decision time and ITI to the inter-trial interval. This reward rate was calculated for congruent and incongruent trials separately, and then summed. The calculation of the reward rate for each combination was based on 20000 simulations from the DDM. The optimal trajectory was defined as the maximum trajectory in the resulting reward rate landscape.

The path analyses were performed by Multivariate Bayesian Linear Mixed Effects modeling using the brms package. The different paths are described by different regression equation which are estimated jointly. We included random slopes for all predictors but did not need random intercepts (for the same reason as above) along with correlation among the random slopes of independent variables between the multiple regression equations. Again, we used weakly informative priors for all beta’s (normal distribution with mean zero and a standard deviation of 2.5). Below, we present the model equations (in the lme4/brms style syntax) for the final path model:

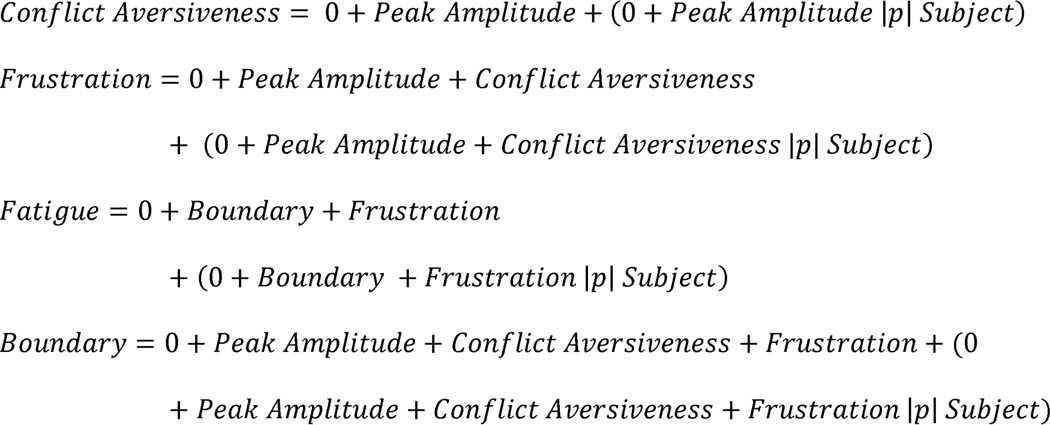

The part of the equation in parentheses is related to the random effects structure. The notation (X |p| Y) denotes that there are random slopes of independent variable X for each grouping of Y (i.e., subject) that are allowed to be correlated (|p|) across the multiple dependent variables. The zeros denote the absence of fixed and random intercepts. The indirect effects are calculated by multiplying the posteriors of the beta’s from two respective paths.

### EEG Data Acquisition & Preprocessing

EEG was recorded from 64 active electrodes (actiCap snap, Brain Products). Impedances were brought below 10 kΩ before starting the experiment. EEG activity was referenced online to the C1 electrode and signals were digitized at a sampling rate of 500 Hz. The EEG recordings were analyzed with MNE (^98^, version 0.24.1) in Python (version 3.9.7). The signal was band-pass filtered between 0.1 and 40 Hz, downsampled to 250 Hz and epoched from -1000 to 1500 ms around Flanker onset. Independent Component Analysis (ICA) was estimated on a more highly filtered version of the data (1 Hz high pass filter) as this leads to more stable solutions of the ICA algorithm and is recommended by MNE. The blink component was automatically identified (and subtracted from the data) as the component with the highest correlation with the two most frontal electrodes (Fp1, Fp2) in which blink activity is most prominent. The automatic selection was manually inspected for correctness (in four subjects, we manually identified a second blink component). Next, an automatic trial rejection procedure (autoreject, see ^99^ for details of the algorithm, version 0.2.2) and a fixed peak-to-peak threshold (150 muV) was used to remove trials with artifacts (*M =* 3.70 %, *SD* = 4.21 %). The resulting epochs were visually inspected (raw epochs and power spectra) to confirm that the procedure worked as expected. Next, we interpolated bad channels (0 channels: N = 27, 1 channel: N = 8, 2 channels: N = 4, 3 channels: N = 6, 4 channels: N = 4) and re-referenced the data to the average of all electrodes.

The Lateralized Readiness Potential (LRP) was calculated for both congruent and incongruent trials using the double difference method ([C3_left_ – C4_left_] – [C3_right_ – C4_right_]) which results in a positive polarity when the correct response hand is activated and a negative polarity when the incorrect response hand is activated. The signal was baseline corrected from -200 to 0 ms and spatially filtered by computing the Laplacian which decreases the effect volume conductivity and produces a more distinct topography. Next, we extracted features from the LRP that could be considered neural surrogates for the DDM parameters. First, we computed the slope of the average LRP between the time of the first incongruent negative peak and the second incongruent peak (“LRP slope”). This timing was chosen to avoid contamination of the signal by impact from the irrelevant dimension (which mainly impacts motor preparation before the first incongruent negative peak). Second, we extracted the positive peak amplitude of the LRPs (“LRP amplitude”) as an approximation to the decision boundary. All neural peak amplitude measures were based on averaging across a window of 50 ms before and after the grand average peak amplitude. Third, the onset latency of the LRPs (“LRP latency”), was estimated using segmented (piecewise) regression (Schwarzenau et al., 1998). Based on visual inspection of the grand average data, we restricted the segmented regression to look for one breakpoint between 0 and 276 ms. Visual inspection indeed revealed that the segmented regression was successful at estimating the onset of motor preparation. Last, we used the (absolute) negative peak amplitude on incongruent trials (“LRP dip”) as a neural proxy for irrelevant capture.

## Data and code availability statement

The data to replicate the results and the associated code will be available upon publication.

## Acknowledgements

We would like to thank Amitai Shenhav and Alan Voodla for valuable comments on a previous draft of the manuscript. W.N., S.B. (G.0660.17N) and L.V. (11H5619N and 1242924N) were supported by the FWO –Research Foundation Flanders. S.B. was supported by an ERC Starting grant (European Union’s Horizon 2020 research and innovation program, Grant agreement 852570). C.G.G. was supported by the Special Research Fund of Ghent University (BOF.GOA.2017.0002.03) and M.R. by the Spanish Ministry of Science and Innovation (PID2019-111187GB-I00). All procedures applied in the present experiment were carried out with adequate understanding and written consent of the subjects and are in accordance with the Declaration of Helsinki.

## Author contributions

L.V.: Conceptualization, Methodology, Data Collection, Formal Analysis, Writing - Original Draft.

S.B.: Conceptualization, Methodology, Supervision, Writing - Review & Editing.

K.D.: Conceptualization, Methodology, Writing - Review & Editing.

I.I.: Conceptualization, Methodology, Writing - Review & Editing.

J.M.G: Data Collection, Writing - Review & Editing.

C.G.G: Supervision, Resources – Equipment, Writing - Review & Editing.

M.R.: Supervision, Resources – Equipment, Writing - Review & Editing.

W.N: Conceptualization, Methodology, Supervision, Writing - Review & Editing.

## Competing interests

The authors have no competing interests to declare.

## Notes

### Competing Interest Statement

The authors have declared no competing interest.

### Summary of Updates

Some textual clarifications have been made in the results and discussion section, based on reviewer feedback

