## Supplementary Materials for "The Temporal Dynamics of Metacognitive Experiences Track Rational Adaptations in Task Performance"

**S1: Behavioral Analyses**

**S2: Fit assessment and parameter recovery of the diffusion modeling data**

**S3: Inter-individual similarity between DDM parameters and neural indices**

**S4: Inter-individual similarity between DDM parameters and metacognitive experiences
S5: Intra-individual similarity between behavioral indices and metacognitive experiences**

**S6: Overview of Path Models**

### S1: Behavioral analyses

Similar to the DDM parameters, we examined the evolution of behavior (RT and Error Rate) with time-on-task across the behavioral and EEG samples (N = 111). We conducted linear mixed effects models predicting RT and Error Rate by time-on-task (i.e., block, 1-18) and congruency (congruent, incongruent). We fitted separate models for linear, logarithmic, quadratic, or cubic effects of Time-on-task and chose the model with the lowest Bayesian Information Criterion (BIC). For the RT measure, the best model was logarithmic (Fig. 2A, left panel). We found an effect of congruency showing the typical interference effect as participants were faster on congruent trials relative incongruent trials, *β* = -0.65, *SE* = 0.009, *t*(1528) = -72.17, *P* < .001. General RT decreased with time-on-task, *β* = -0.123, *SE* = 0.026, *t*(44) = -4.78, *P* < .001 and this effect interacted with congruency, *β* = 0.068, *SE* = 0.009, *t*(1528) = -7.50, *P* < .001, as the decrease was stronger for incongruent trials, *b* = -8.43, *SE* = 1.20, relative to congruent trials, *b* = -2.42, *SE* = 1.20, thus resulting in a decreasing interference effect with time-on-task.

In the Error Rate measure, the best model was again logarithmic (Fig 2A, right panel). We found the typical interference effect as participants made more errors on incongruent trials relative to congruent trials, *β* = -0.693, *SE* = 0.014, *t*(1528) = -48.984, *P* < .001. Error Rate did not change with time-on-task, *β* = 0.011, *SE* = 0.027, *t*(44) = 0.424, *P* = 0.674. However, congruency interacted with time-on-task, *β* = 0.032, *SE* = 0.014, *t*(1528) = 2.290, *P* = 0.022, as the Error Rate on congruent trials tended to increase, *b* = 0.004, *SE* = 0.003, whereas it tended to decrease on incongruent trials, *b* = -0.002, *SE* = 0.003, thus again resulting in a decreased interference effect over time-on-task.

### S2: Fit assessment and parameter recovery of the diffusion modeling data

We assessed whether the diffusion model captured the behavioral data by inspecting the observed versus predicted RT quantiles and conditional accuracy function for congruent and incongruent trials separately. We found that the diffusion model fit the human data well (Supplementary Fig. 1A). Next, we simulated 100 datasets from our drift diffusion model. Each dataset contained 260 trials with 50 % congruent trials and 50% incongruent trials (i.e., the exact same properties as one block of experimental data). For each simulation, a random set of parameters were drawn from a uniform distribution within the parameter boundaries as described in the Methods section. Next, we used the same fitting procedure as for the experimental data to check whether the true parameter values would be recoverable. Supplementary Fig. 1B shows that we were indeed able to adequately recover the true parameter values (all *R* > .75) given the number of trials used for fitting the data.

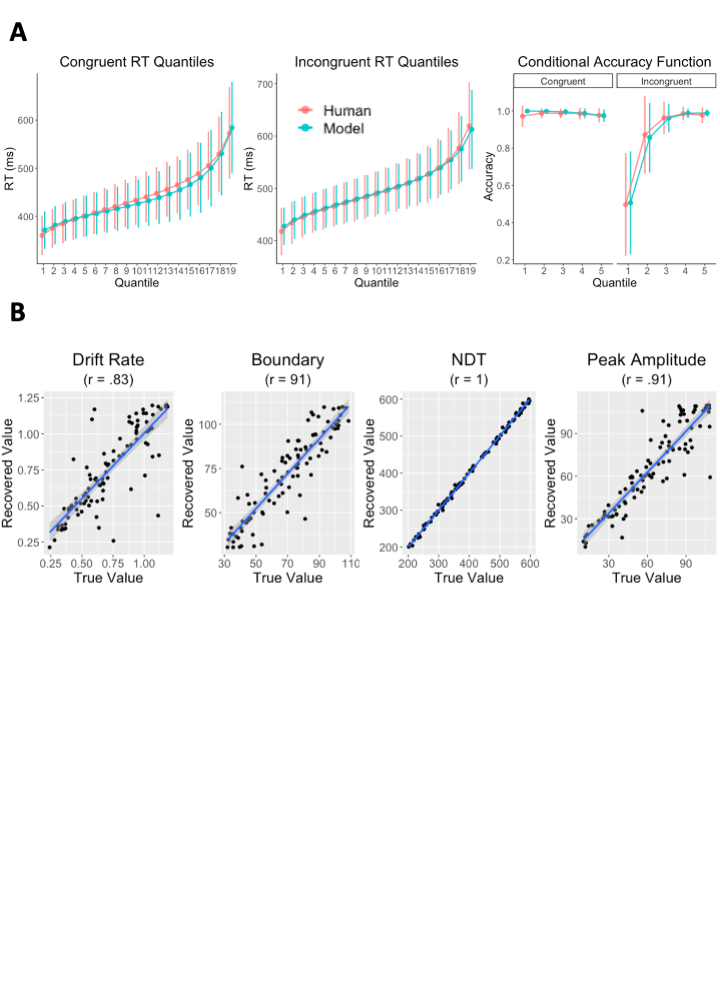
 **Supplementary Fig. 1: Fit assessment and parameter recovery. (A)** Reaction time and accuracy of human observations versus model predictions. **(B)** Correlations between generating (true) and recovered parameter values. NDT = Non-decision Time.

### S3: Inter-individual similarity between DDM parameters and neural indices

In addition to the intra-individual relationships between DDM parameters and neural indices reported in the main text, we also investigated the inter-individual relationships using Canonical Correlation Analysis (CCA). CCA is a dimensionality reduction technique that identifies novel linear projections or “dimensions” of two datasets that maximizes the correlation between them^1,2^. CCA was performed on the standardized individual intercepts and first-degree slopes of all DDM parameters on the one hand (“DDM domain”; 10 variables) and all neural indices on the other hand (“EEG domain”, 8 variables).

We found that the dimensionality of the subspace that maximally relates the two domains is sufficiently described as unidimensional, *R* = .806, *Wilks’ Lambda (for dimension 1 through 8)* = 0.035, *F_approx._*(80, 179.81) = 1.561, *P* = 0.008. For the DDM domain, we found that intercepts and slopes of boundary, peak amplitude and peak-to-bound ratio contributed most strongly to this dimension (Supplementary Fig. 2A), showing a strong focus on the degree of and changes in model-based irrelevant capture. For the EEG domain, we found that intercepts and slopes of LRP dip and slopes of LRP slope contributed most strongly to this dimension (Supplementary Fig. 2B), showing a strong focus on the degree of and changes in neural-based irrelevant capture. Follow-up multiple regressions predicting each of the three strongest contributors from the EEG domain by the strongest contributors from the DDM domain (excluding intercepts and slopes of peak amplitude to avoid multicollinearity with peak-to-bound ratio and boundary) revealed that intercepts and slopes of LRP dip where most strongly related to intercepts and slopes of peak-to-bound ratio respectively (intercepts LRP dip – intercepts peak-to-bound ratio: *β* = 0.535, *SE* = 0.193, *t*(39) = 2.75, *P* = .008, slopes LRP dip – slopes peak-to-bound ratio: *β* = 0.397, *SE* = 0.150, *t*(39) = 2.64, *P* = .012). On the other hand, slopes of LRP slope were specifically related to slopes of drift rate, *β* = 0.472, *SE* = 0.186, *t*(39) = 2.54, *P* = .015.

In sum, we can conclude that this canonical correlation especially highlights the similar nature of how irrelevant capture and evidence accumulation are represented by the model parameters and the neural indices.

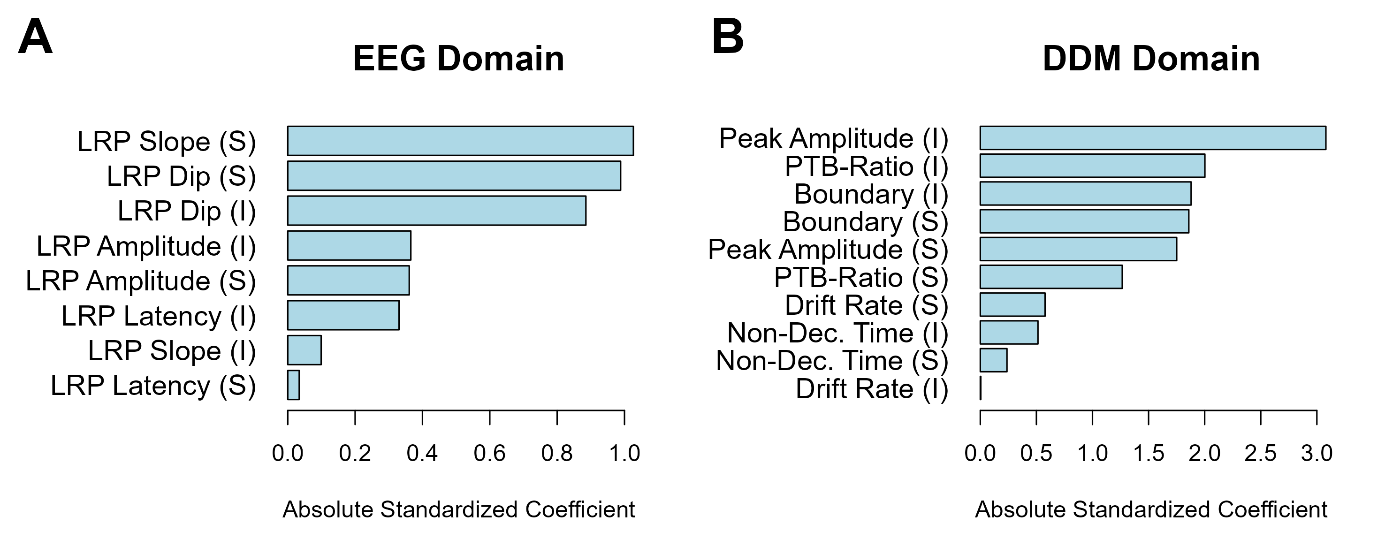

**Supplementary Fig. 2: Coefficients from the Canonical Correlation Analysis (CCA) between the DDM and EEG domain. (A)** Contributions of the variables to the first dimension for the EEG Domain. **(B)** Contributions of the variables to the first dimension for the DDM domain. (I) = Intercept, (S) = Slope, LRP = Lateralized Readiness Potential, PTB = Peak-to-bound ratio.

### S4: Inter-individual similarity between DDM parameters and metacognitive experiences

For investigation of the similarity between DDM parameters and metacognitive experiences at the inter-individual level, we again turned to CCA on the individual intercepts and first-degree slopes of the DDM parameters (10 variables) and Metacognitive experiences (10 variables). We again found that a unidimensional representation was sufficient, *R* = .671, *Wilks’ Lambda (for dimension 1 through 10)* = 0.204, *F_approx._*(100, 663.39) = 1.648, *P* < 0.001. For the Metacognitive domain, we found a clear domination of this first dimension by slopes in conflict aversiveness (Supplementary Fig. 3A). For the DDM domain, we found that intercepts of peak amplitude and slopes of peak-to-bound ratio (and to a lesser degree intercepts of non-decision time and boundary) contributed most strongly to this dimension (Supplementary Fig. 3B). As a follow up, we performed a multiple regression predicting slopes of conflict aversiveness by the strongest contributors of the DDM domain. This revealed a strong relationship between slopes of peak-to-bound ratio and slopes of conflict aversiveness, *β* = 0.402, *SE* = 0.091, *t*(106) = 4.37, *P* < .001. This finding is interesting, as we did not observe changes in conflict aversiveness with time-on-task averaged across the sample. However, these results suggest that there are meaningful subgroups with respect to conflict aversiveness, based on how irrelevant capture evolved throughout the experiment (i.e., peak-to-bound ratio). Linear mixed effects modeling of the conflict aversiveness time-on-task effects by the peak-to-bound ratio slopes showed that when people were not able to decrease irrelevant capture during the experiment (a stable trajectory of peak-to-bound ratio), they showed more conflict aversion (Supplementary Fig 3C). On the other hand, when people were able to strongly decrease the degree of irrelevant capture, they showed less conflict aversion.

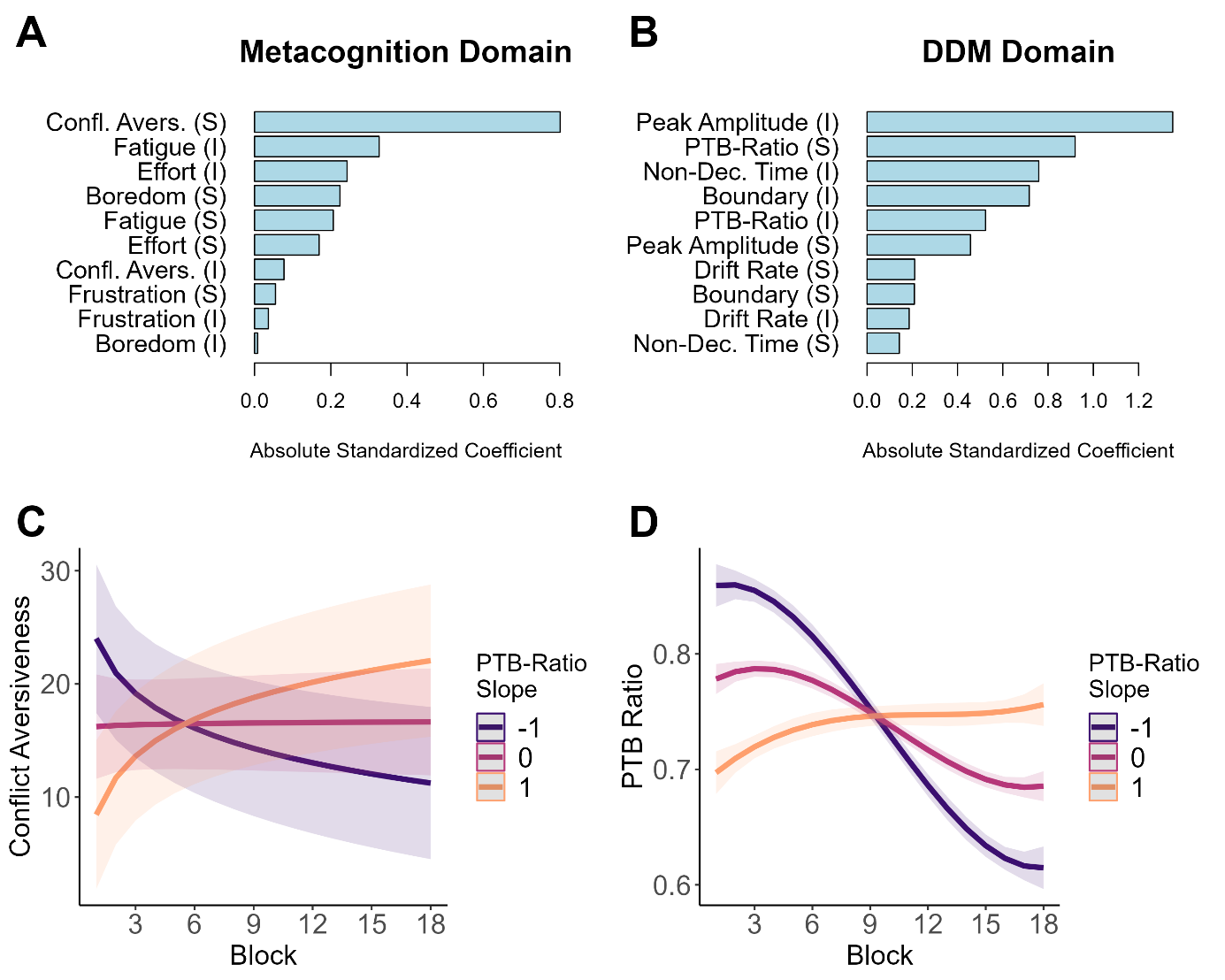

**Supplementary Fig. 3: Coefficients from the Canonical Correlation Analysis (CCA) between the Metacognition and DDM domain. (A)** Contributions of the variables to the first dimension for the metacognition domain. **(B)** Contributions of the variables to the first dimension for the DDM domain. **(C)** Trajectories of conflict aversiveness for varying individual slopes of peak-to-bound ratio. **(D)** Trajectories of peak-to-bound ratio for varying levels of individual slopes of peak-to-bound ratio (to demonstrate the different trajectories of individuals with varying levels of peak-to-bound ratio slopes). (I) = Intercept, (S) = Slope, Confl. Avers. = Conflict aversiveness, LRP = Lateralized Readiness Potential, PTB = Peak-to-bound ratio.

### S5: Intra-individual similarity between behavioral indices and metacognitive experiences

For investigation of the similarity between behavioral indices and metacognitive experiences at the intra-individual level, we conducted a Bayesian Multivariate Linear Mixed Effects Model to predict local fluctuations of the behavioral indices (average RT and accuracy across congruency levels and the congruency effects [CE] in RT and accuracy) by local fluctuations of the metacognitive experiences (Supplementary Fig. 4). This analysis revealed that fatigue and frustration were negatively related to accuracy, indicating that blocks with higher levels of fatigue and frustration corresponded to lower levels of accuracy. For frustration, we found additional positive relationships between the congruency effects (CE) in RT and accuracy, with higher levels of frustration corresponding to larger congruency effects (a similar pattern was observed for the experience of conflict aversiveness). This shows that, at the behavioral level, fatigue and frustration predominantly align with "negative" changes in behavior, such as decreased RT or increased interference. Notably, lower RTs (“better” performance) were specifically linked to effort, indicating that performance improvement was primarily driven by the exertion of cognitive effort. These findings resonate with our interpretation that fatigue and frustration are not indicative of successful behavioral adaptations per se, but rather manifestations of the ongoing, cognitively taxing process of constant behavioral adjustment. Rather than signaling success or increased task ease, they reflect the output of this effortful process as it is decreasing subjects’ cognitive capacity.

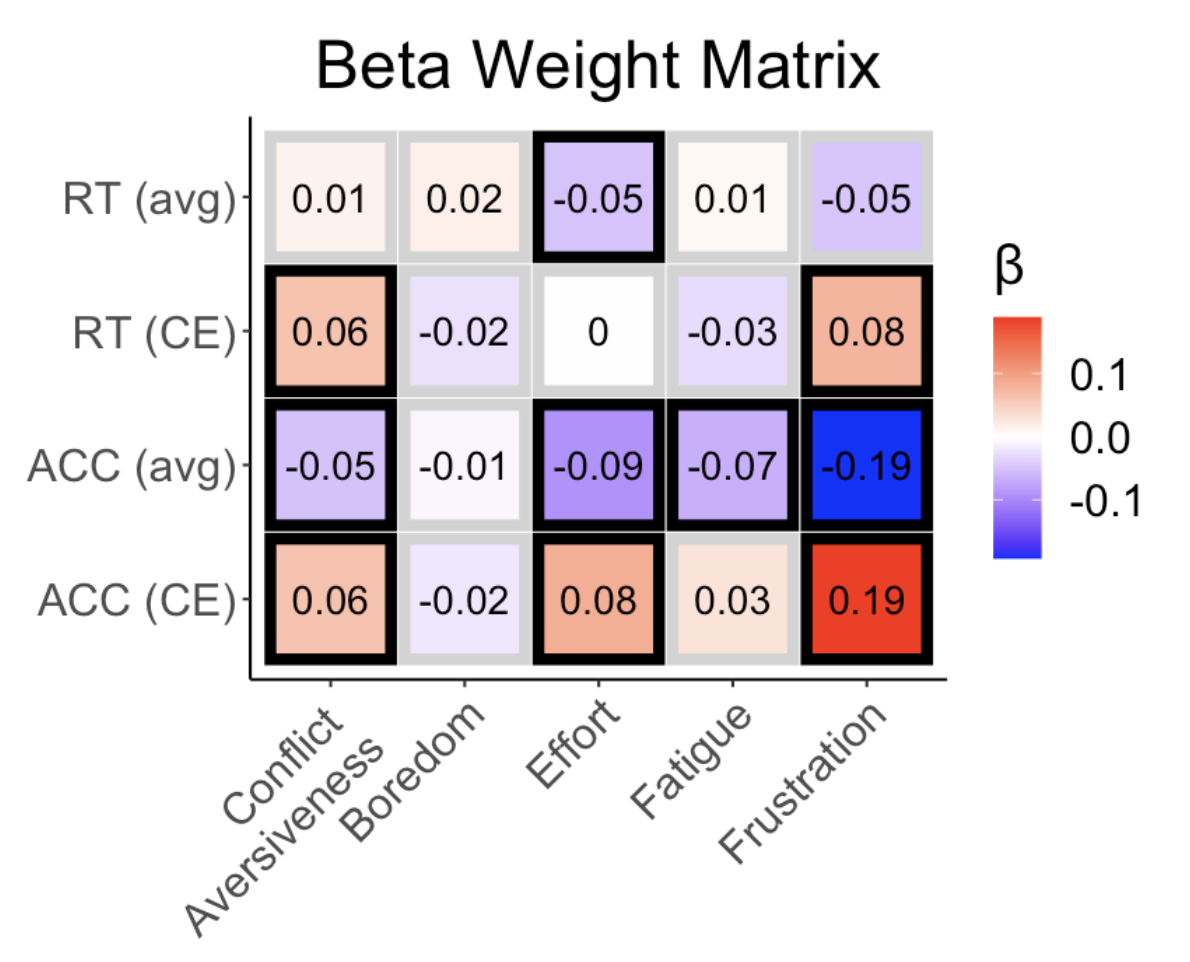

**Supplementary Fig. 4:** Beta weight matrix for metacognitive experiences and raw behavioral indices. Avg = Average (across congruency conditions). CE = congruency effect (also known as interference effect). Black boxes denote coefficients where the 95 % HDI does not contain zero.

### S6: Overview of path models

The following Supplementary Figures (5A-E) portray the model building steps followed building the final path model. In a first model, we started with a parallel mediation model where all metacognitive experiences are allowed to mediate the relationship between peak amplitude and boundary (Supplementary Fig. 5A). No significant mediators were found. Because we previously found that fatigue was related to the decision boundary, we test in a second model (Supplementary Fig 5B) whether boundary predicts fatigue, rather than fatigue mediating the relationship between peak amplitude, which we found to be the case. Next, in a third model (Supplementary Fig 5C), we investigated if there was a path connecting the peak amplitude - conflict aversiveness and the boundary - effort/frustration relationships. Again, no significant mediators found, but a path opened over conflict aversiveness and effort. In a fourth model Supplementary Fig. 5D), we found evidence for the indirect path (over conflict aversiveness, effort and frustration) that opened in the previous model. Finally, in model 5 (the final model reported in the manuscript; Supplementary Fig. 5E), we investigated if there was a relationship between frustration and fatigue and whether boundary would qualify as a mediator for that relationship, which we indeed found.

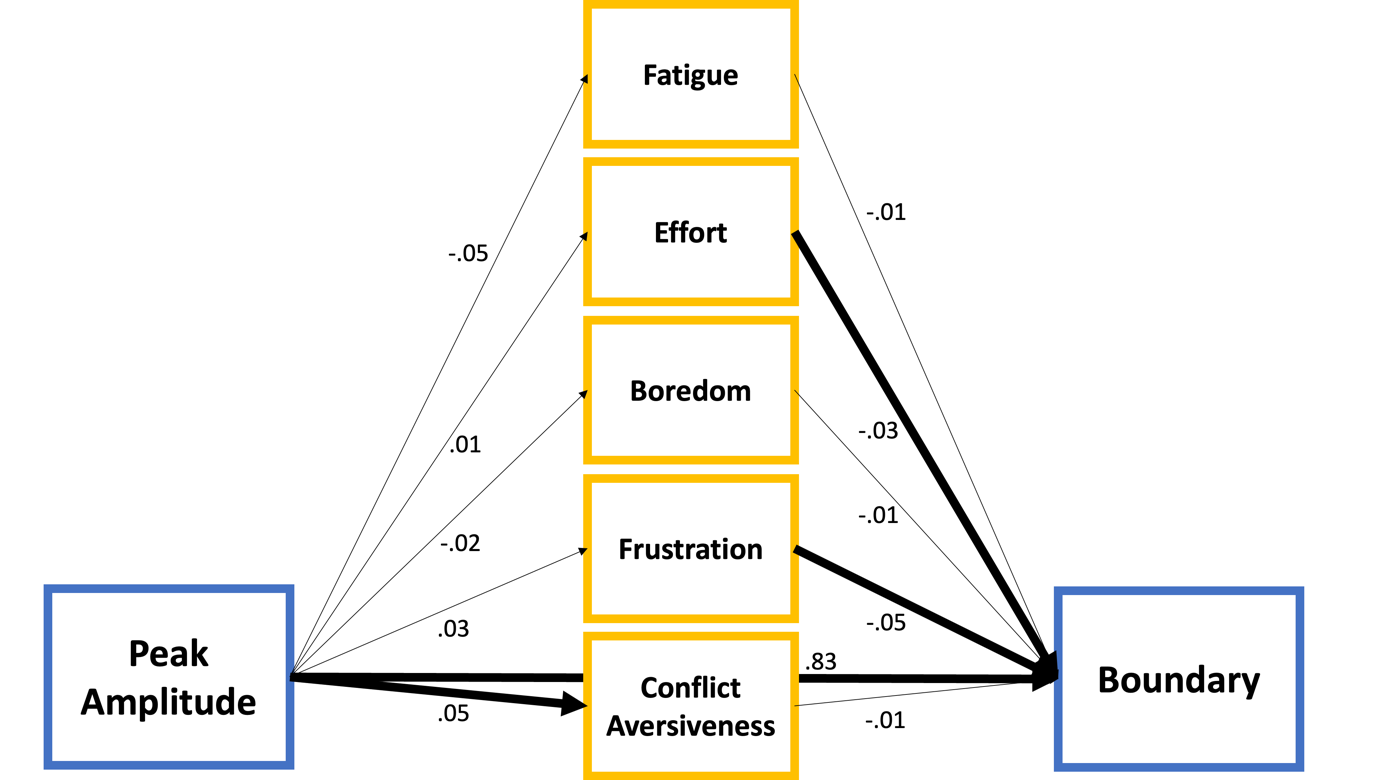

**Supplementary Fig. 5A:** Path Model 1. Significant relationships (for which the 95% CI does not include zero) have dark arrows. Blue relates to task performance parameters, yellow to metacognitive experiences.

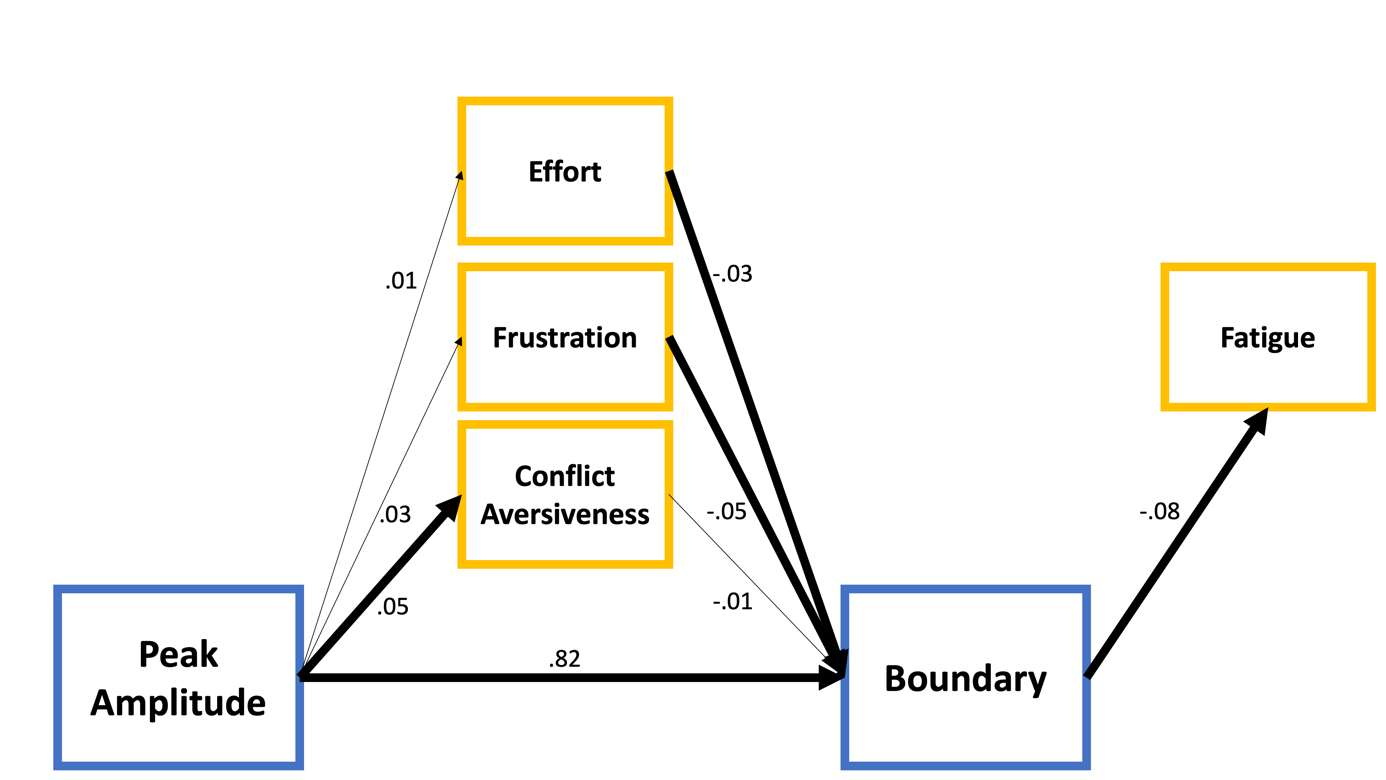

**Supplementary Fig. 5B:** Path Model 2. Significant relationships (for which the 95% CI does not include zero) have dark arrows. Blue relates to task performance parameters, yellow to metacognitive experiences.

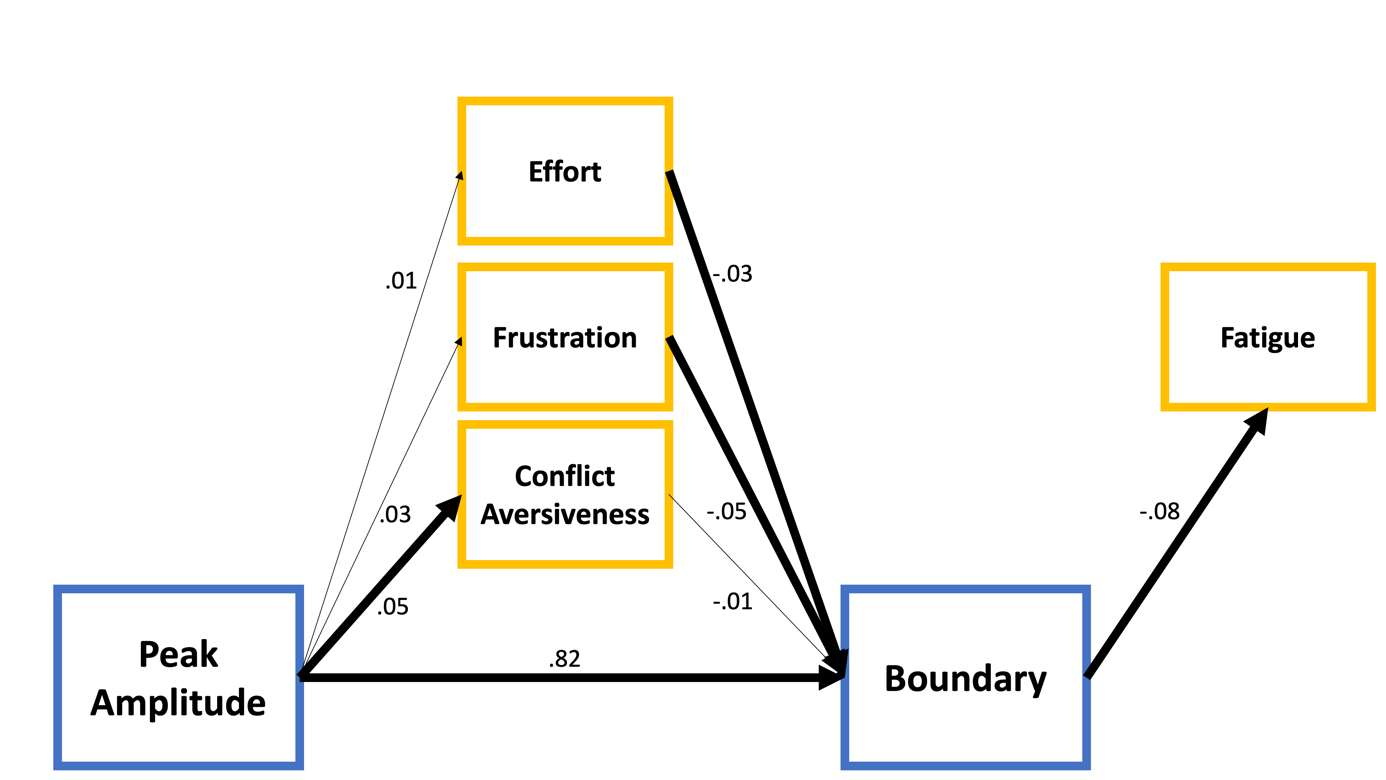

**Supplementary Fig. 5C:** Path Model 3. Significant relationships (for which the 95% CI does not include zero) have dark arrows. Blue relates to task performance parameters, yellow to metacognitive experiences.

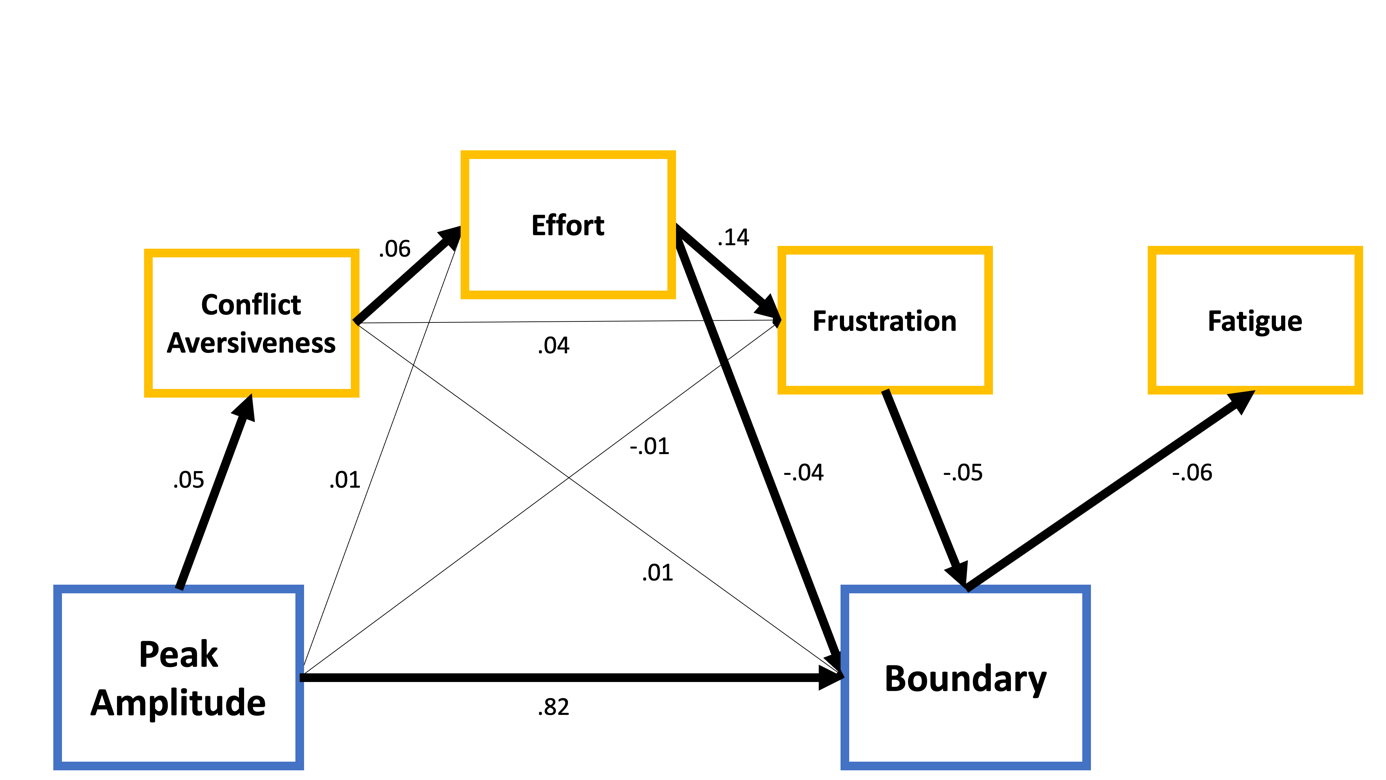

**Supplementary Fig. 5D:** Path Model 4. Significant relationships (for which the 95% CI does not include zero) have dark arrows. Blue relates to task performance parameters, yellow to metacognitive experiences.

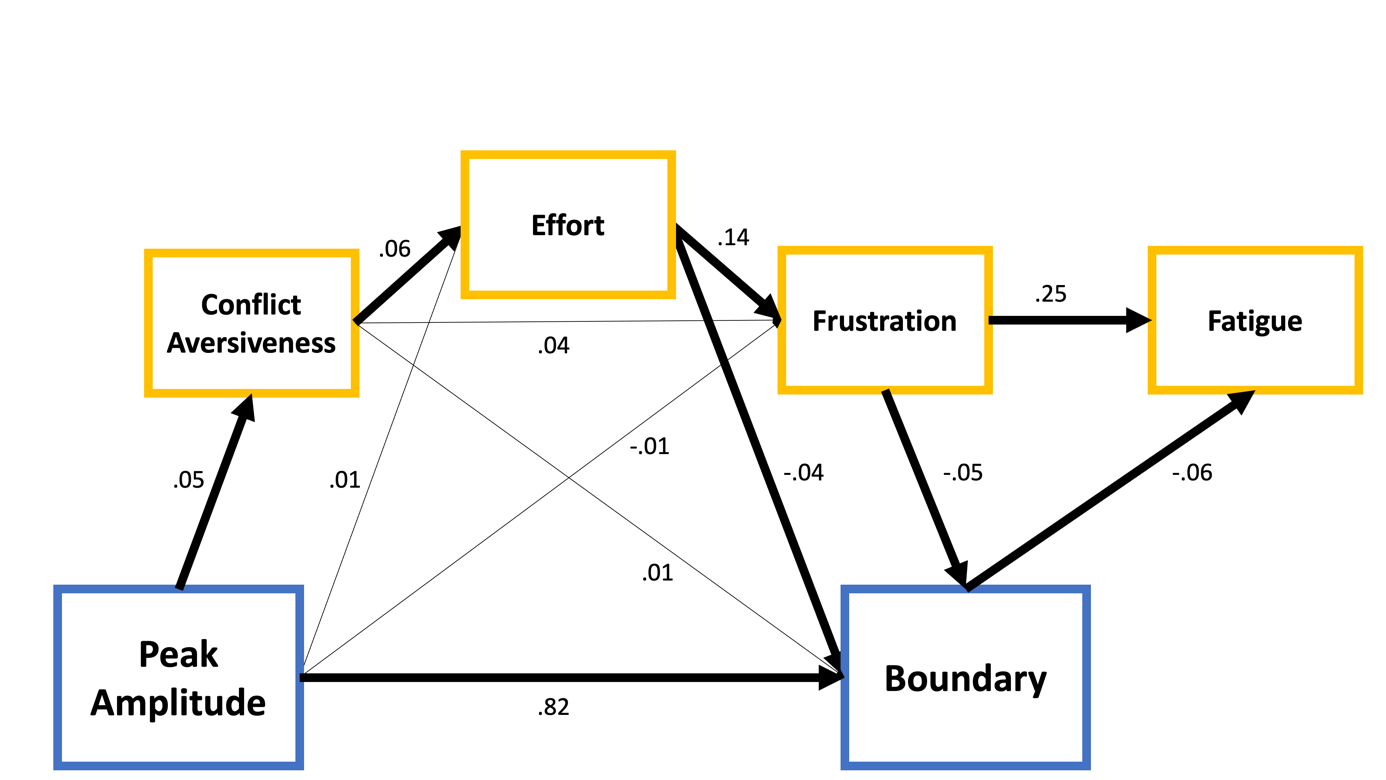

**Supplementary Fig. 5E:** Path Model 5. Significant relationships (for which the 95% CI does not include zero) have dark arrows. Blue relates to task performance parameters, yellow to metacognitive experiences.
